# Local interactions lead to spatially correlated gene expression levels in bacterial groups

**DOI:** 10.1101/109991

**Authors:** Simon van Vliet, Alma Dal Co, Annina R. Winkler, Stefanie Spriewald, Bärbel Stecher, Martin Ackermann

**Affiliations:** Institute of Biogeochemistry and Pollutant Dynamics, Department of Environmental Systems Science, ETH Zurich, Zurich, Switzerland; Department of Environmental Microbiology, Eawag, Dübendorf, Switzerland; Max-von-Pettenkofer Institute, LMU Munich, Munich, Germany; German Center for Infection Research (DZIF), partner site LMU Munich, Munich, Germany

## Abstract

Many bacteria live in spatially structured assemblies where the microenvironment of a cell is shaped by the activities of its neighbors. Bacteria regulate their gene expression based on the inferred state of the environment. This raises the question whether the phenotypes of neighboring cells can become correlated through interactions via the shared microenvironment. Here, we addressed this question by following gene expression dynamics in *Escherichia coli* microcolonies. We observed strong spatial correlations in the expression dynamics for pathways involved in toxin production, SOS-stress response, and metabolism. These correlations can partly be explained by a combination of shared lineage history and spatial gradients in the colony. Interestingly, we also found evidence for cell-cell interactions in SOS-stress response, methionine biosynthesis and overall metabolic activity. Together our data suggests that intercellular feedbacks can couple the phenotypes of neighboring cells, raising the question whether gene-regulatory networks have evolved to spatially organize biological functions.

## Introduction

Many bacteria do not live in isolation, instead they are members of larger communities (Claessen et al. 2014). The microenvironment in these communities is partly determined by abiotic conditions, but it is also affected by the activities of a cell’s neighbors (Flemming et al. 2016; Stewart and Franklin 2008). At the same time, cells adjust their phenotype by regulating gene expression based on the inferred state of the local microenvironment. The phenotype of a cell is thus likely influenced by its location in the community and by the identity and the activities of neighboring cells.

The functionality of the community as a whole depends on the combined activities of all of its members. Being part of a group can allow cells to specialize in performing different tasks (Kolter, Vlamakis, and van Gestel 2015). Such interactions between different cell types can lead to new or improved functionality that goes beyond the sum of the activities of the individual cells (Claessen et al. 2014; van Vliet and Ackermann 2015). In multispecies biofilms most of this specialization is the result of genetic differences between member species. However, specialization can likewise occur in single species communities as the result of phenotypic variation among cells (Kolter, Vlamakis, and van Gestel 2015; Ackermann 2015; Ackermann et al. 2008). One well studied example is in clonal *Bacillus subtilis* biofilms, where functionality depends on interactions between multiple different cell types (Lopez and Kolter 2010; Kolter, Vlamakis, and van Gestel 2015).

Interactions between different cell types can be sensitive to the spatial arrangements of the different types (Liu et al. 2016; Nadell, Drescher, and Foster 2016). For example, the division of labor between nitrogen fixing and photosynthetic cells in multicellular cyanobacteria is likely more efficient due to the regular spacing of nitrogen fixing cells along the filaments (Muro-Pastor and Hess 2012). Furthermore, recent work in *B. subtilis* colonies directly linked functionality at the group level to the spatial arrangement of two cell types that perform complementary functions (van Gestel, Vlamakis, and Kolter 2015). More generally, we expect that correlations in the phenotypes of neighboring cells can be beneficial for a large number of activities (Ross-Gillespie and Kümmerli 2014). Positive correlations (i.e. neighbors having similar phenotypes) can allow cells to coordinate their activities. This can be of benefit in the production of secreted effectors, by allowing a sufficient build-up in local effector concentrations. Negative correlations (i.e. neighbors having different phenotypes) can facilitate division of labor between cells. This can be of benefit in the context of anabolic pathways: cells can benefit from economies of scale by specializing on the biosynthesis of a subset of metabolites, while exchanging end products with neighbors specializing on complementary pathways (Johnson et al. 2012; Guantes et al. 2015).

Spatial correlations in phenotypes can be the result of several processes. In previous work it was found that several phenotypes can be epigenetically inherited (Robert et al. 2010; Veening et al. 2008; Veening, Smits, and Kuipers 2008; Hormoz, Desprat, and Shraiman 2015; Julou et al. 2013). Such epigenetic inheritance leads to positive correlations between the phenotypes of closely related cells (e.g. between sisters or cousins, i.e., cells that are only separated by one or two cell divisions (Hormoz, Desprat, and Shraiman 2015)). As neighboring cells tend to be highly related, epigenetic inheritance can lead to positive spatial correlations in phenotypes. Furthermore, neighboring cells are exposed to a similar combination of environmental gradients (Flemming et al. 2016; Stewart and Franklin 2008). A common gene regulatory response to these gradients can thus result in positive correlations in phenotypes. Finally, local interactions (i.e. intercellular feedbacks) can lead to a coupling in expression levels between neighbors (Risser, Wong, and Meeks 2012; Bassler and Losick 2006). These intercellular feedbacks can give rise to either positive or negative correlations in phenotypes. Spatial correlations in phenotypes will depend on the combined effects of epigenetic inheritance, spatial gradients, and intercellular feedbacks. However, so far these processes have mostly been studied in isolation. At the moment we thus lack a quantitative understanding of the relative contributions of these processes to spatial correlations in phenotype.

Here we address the following question: to what extent are cellular activities correlated between neighboring cells? We are especially interested in the role of intercellular feedbacks between neighbors. As a model system we used two-dimensional microcolonies of *Escherichia coli.* We followed the growth of a microcolony using time-lapse microscopy while tracking spatiotemporal gene expression patterns using transcriptional reporters. Subsequently, we quantified spatial correlations in gene expression and developed a novel statistical approach to disentangle the effects of shared linage history, spatial gradients, and local interactions. We found strong spatial correlations for all studied pathways, which we could attribute to the effects of shared lineage history and, for some pathways, the effects of spatial gradients and local interactions.

We studied two groups of pathways for which we expected to find spatial correlations of opposite signs. As a model for secreted effector molecules, where we expected positive correlations, we chose the bacteriocin colicin Ib. In natural systems the extra cellular concentration of bacteriocin needs to reach a threshold concentration to inhibit the growth of nearby sensitive cells (Cascales et al. 2007). If neighboring cells coordinated their expression dynamics, it would be easier for them to reach this threshold concentration. We thus hypothesized that colicin Ib expression levels should be positively correlated between neighboring cells.

As a model for anabolic pathways, where we expected negative correlations, we chose three pathways involved in amino acid biosynthesis. Previous work using genetic consortia of complementary aminoacids autotrophs has shown that many amino acids can be exchanged through the environment (Mee et al. 2014; Wintermute and Silver 2010). Furthermore, amino acid production costs appear to show economies of scale: the cost of producing an extra amino acid decreases with increasing production levels (Pande et al. 2013). The community can thus grow faster if its members specialize on the production of complementary subset of amino-acids and exchange surpluses with each other. Indeed, it was found that several genetic consortia of cross-feeding strains (i.e. complementary strains that cannot produce one amino acid but overproduce another) could outcompete a wildtype ancestor (Pande et al. 2013). This shows that a division of labor strategy can increase population growth rates when cells are genetic specialists. We hypothesized that a similar benefit could be obtained if genetically identical cells phenotypically specialize on the production of different amino acids. Such phenotypic specialization could be obtained if random fluctuations in amino acid production rates (e.g. due gene expression noise (Elowitz et al. 2002; Ozbudak et al. 2002; Kaern et al. 2005)) are amplified by intercellular feedback loops. Furthermore, we hypothesized that cells would arrange their activities in space to optimize the efficiency of the division of labor. Specifically, we expected that neighboring cells would specialize on different pathways in order to optimize the exchange of amino acids. We thus expected to observe negative correlations (i.e. dissimilarity) in the expression levels of amino acid synthesis pathways between neighboring cells.

## Results

To test our hypotheses, we followed the spatiotemporal gene expression dynamics in microcolonies of *E.coli*. Gene expression was quantified using plasmid-based green fluorescent protein (GFP) transcriptional reporters. The mean fluorescent intensity of the transcriptional reporter is approximately proportional to the concentration of GFP and hence a proxy for the concentration of the protein encoded by the gene of interest. We thus refer to this quantity as *protein level*. We furthermore quantified the rate of change in the total fluorescent intensity over time, which is a proxy for the *promoter activity* (see *Methods*, (Kiviet et al. 2014)). Cells were inoculated at low density on agar pads and the growth of a microcolony founded by a single cell was followed over time using time-lapse microscopy. We quantitatively analyzed the observed spatiotemporal gene expression patterns using a newly developed statistical approach. We will first describe the dynamics observed for colicin Ib, while simultaneously introducing our analysis methods. Subsequently, we will apply the same methods to analyze the dynamics for the amino acid synthesis pathways.

### Neighboring cells have similar protein levels of colicin Ib

First we investigated the expression patterns of the bacteriocin colicin Ib (*cib*), which is a pore-forming toxin found in natural isolates of *E. coli* and *Salmonella* (Riley and Wertz 2002; Cascales et al. 2007). *cib* transcription is co-repressed by the binding of LexA to the SOS-box (SOS DNA repair response) and by the binding of Fe^2+^-Fur complex to the iron box (Ferric uptake regulation) (Cascales et al. 2007; Nedialkova, Denzler, Koeppel, Diehl, Ring, Wille, Gerlach, and Stecher 2014a). *cib* transcriptional repression can be partly relieved by activation of the SOS-response in response to DNA damage, or by a shortage in ferrous iron (Fe^2+^) (Spriewald et al. 2015; Nedialkova, Denzler, Koeppel, Diehl, Ring, Wille, Gerlach, and Stecher 2014a). Here, we relived repression by Fur by chelating free iron in the medium. We did not induce SOS response, so any *cib* expression is likely the result of SOS induction due to spontaneous occurring DNA damage (Pennington and Rosenberg 2007).

Colicin Ib protein levels varied strongly between cells in a microcolony (median coefficient of variation=0.19, *n*=8, Figure 1A). This is consistent with previous reports of high variation in colicin expression levels (Silander et al. 2012). It is important to note that we do not expect any genetic variation between cells in the microcolony: when a microcolony is founded by a single cell there is a 88% chance that no mutations occur after 7 generations (assuming a mutation rate of 10^-3^, per genome, per generation (Lee et al. 2012)). Likewise, we do not expect abiotic variation in the agar pads: diffusion should equalize any inhomogeneities across the colony within seconds (e.g. a molecule with a diffusion coefficient similar to that of glucose (D~600 μm^2^/s) diffuses across a microcolony (~13μm) in approximately 0.07 seconds). Instead, gene expression noise is likely an important factor leading to the observed phenotypic variation (Elowitz et al. 2002; Ozbudak et al. 2002; Kaern et al. 2005). Moreover, the combined metabolic activities of cells in the colony can give rise to emergent gradients in nutrients and other excreted metabolites. These emergent gradients can in turn also contribute to phenotypic variation.

**Figure 1.**
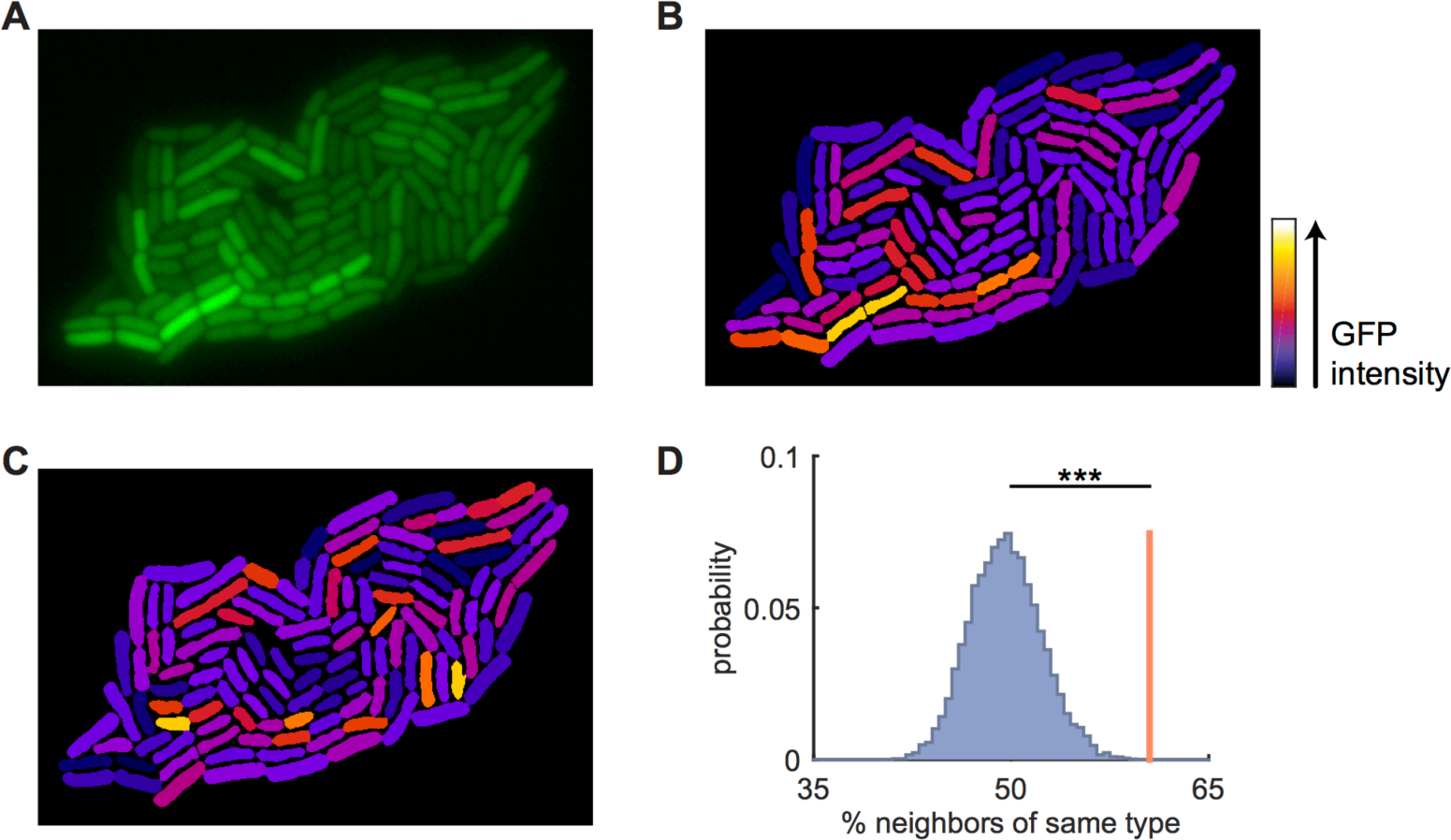
Neighboring cells have similar expression levels of colicin Ib. **A)**Fluorescence image of an *E. coli* microcolony with GFP transcriptional reporter for colicin Ib (*cib*). Colicin expression was induced by chelating free iron in the medium, but no DNA damaging agent was added. **B)** Reconstructed image of the colony shown in A: cell shapes obtained from cell segmentation are uniformly colored based on their mean corrected intensity (see Figure 1–Figure Supplement 1). Note how neighboring cells tend to have similar intensities. **C)** Same as in B, but fluorescence intensities are randomly permuted among the cells. Note that the similarity between neighboring cells has been reduced compared to B. **D)** Quantification of spatial correlations in expression levels: cells are grouped into two clusters (intensity above or below median intensity). Each cell is compared to all its neighbors and the percentage of neighbors that is of the same cell type is calculated (red line). This procedure is repeated 10^4^ times after randomly permuting the intensities among the cells (blue distribution). The observed similarity is significantly higher for the true data compared to the randomized data (p<1·10^-4^, randomization test).

Visual inspection suggested that Colicin Ib levels were non-randomly distributed in the colony. Rather, neighbors appeared to be similar to each other (Figure 1A). To quantitatively investigate the expression patterns, we first corrected the fluorescent images for optical artifacts (Figure 1B,–Figure Supplement 1, see *Methods*). Subsequently, we quantified the spatial correlation in Colicin Ib levels using a randomization test. We found that neighboring cells are significantly more similar to each other than can be expected by chance (Figure 1D, p=10^-4^, see *Methods*). We observed similar results for an additional 8 replicate microcolonies (Figure 1–Figure Supplement 2). Consistent with our hypothesis, we thus consistently observed positive spatial correlation in Colicin Ib levels.

Two main factors could contribute to the observed positive spatial correlation in Colicin Ib levels: shared lineage history and spatial proximity. If a cell’s phenotype is epigenetically inherited (e.g. due to protein inheritance) this will cause similarity in protein levels between closely related cells (e.g. between sisters or cousins, (Veening, Smits, and Kuipers 2008; Hormoz, Desprat, and Shraiman 2015)). As closely related cells also tend to lie close by in space, epigenetic inheritance could lead to spatial correlations in expression levels. Spatial proximity by itself, i.e. independent of the effects of lineage history, could also lead to spatial correlations in expression levels. In this case neighboring cells would be more similar to each other than could be expected based on their relatedness. This could be either due to local cell-cell interactions or due to cells sharing the same microenvironment. As lineage history and spatial proximity tend to be strongly correlated, we developed a statistical method to disentangle these two effects.

### Shared lineage history leads to spatial correlations in colicin Ib expression dynamics

To identify the causes of spatial correlations in Colicin Ib protein levels we followed the growth of microcolonies using time-lapse microscopy and reconstructed the full, spatially resolved, lineage trees (Figure 2A). These lineage trees contain complete information on the locations of cells, their phenotypes, and their relatedness. These lineage tree thus allowed us to disentangle the effects of shared lineage history and spatial proximity on spatial correlations in expression levels.

**Figure 2.**
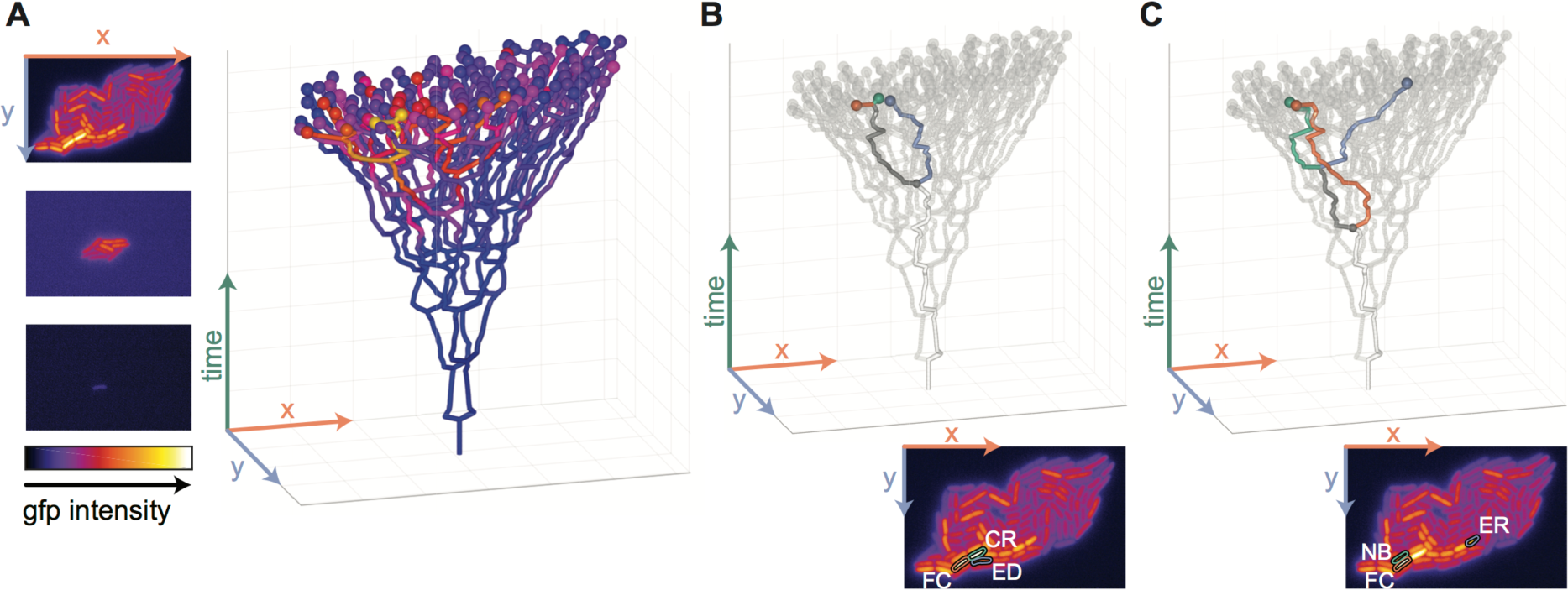
Reconstructing lineage trees to disentangle the effects of space and relatedness. **A)** Left: frames from a time-lapse movie of a growing microcolony with a GFP reporter for *cib*. The images show GFP intensities using a heatmap representation for t=0,3,6h. Right: reconstructed lineage tree. Cells are plotted as a function of location (horizontal plane) and time (vertical axis). Branching points in the lineage tree mark cell division events. The spheres at the tip of the tree represent cells at the final time point with their color indicating the Colicin Ib level of the cell. **B)** Statistical test to quantify the effect of shared lineage history on similarity in expression levels. A focal cell (FC, red) is compared with its closest relative (CR, green) and with an *equidistant cell* (ED, blue), which is a cell that has the same distance to the focal cell as the closest relative, but that is less related. **C)** Statistical test to quantify the effect of spatial proximity on similarity in expression levels. A focal cell (FC, red) is compared with one of its neighbors (NB, green) and with an *equally-related cell* (ER, blue), which is a cell that has the same relatedness to the focal cell as the neighbor, but that is further away in space. **B,C)** The insets at the bottom show the positions of these cells in the GFP image for the last time point (see panel A).

First, we developed a test for the effect of shared lineage history on spatial correlations in phenotype. We could disentangle the effect of relatedness from the effect of spatial proximity by analyzing a group of cells that differ in their relatedness, but that are identical in their spatial arrangement. Specifically, we compared three cells: a focal cell, its closest relative (e.g. its sister), and an *equidistant cell*, i.e. a cell that shares the same spatial relation with the focal cell as the closest relative, but that is less related (Figure 2B). We then compared the phenotypic difference between the focal cell and the *equidistant cell* (*δ_ED_*) with the phenotypic difference between the focal cell and its closest relative (*δ_CR_*). The effect of shared lineage history was quantified by the ratio of these two quantities (i.e. *δ_ED_*/*δ_CR_*). Values larger than 1 indicate that shared lineage history leads to similarity in phenotype (i.e. positive correlations), while values smaller than 1 indicate that shared lineage history leads to dissimilarity in phenotype (i.e. negative correlations).

We found that shared lineage history leads to positive correlations in Colicin Ib levels: a cell is much more similar to its closest relative than to an equally distant (but less related) cell. Specifically, the phenotypic difference between a focal cell and the *equidistant cell* is on average 5.8 times higher than the phenotypic difference between the focal cell and its closest relative (Figure 3A, 〈*δ_ED_*/*δ_CR_*〉=5.8, p<1·10^−5^, t-test, *n*=9). Additionally, we found that closely related cells are also similar with respect to their *cib* promoter activity (Figure 3A, 〈*δ_ED_*/*δ_CR_*〉=1.8, p=0.002). While the similarity in protein level is to be expected due to protein inheritance, the similarity in promoter activity shows that closely related cells are also similar in their current activities.

**Figure 3.**
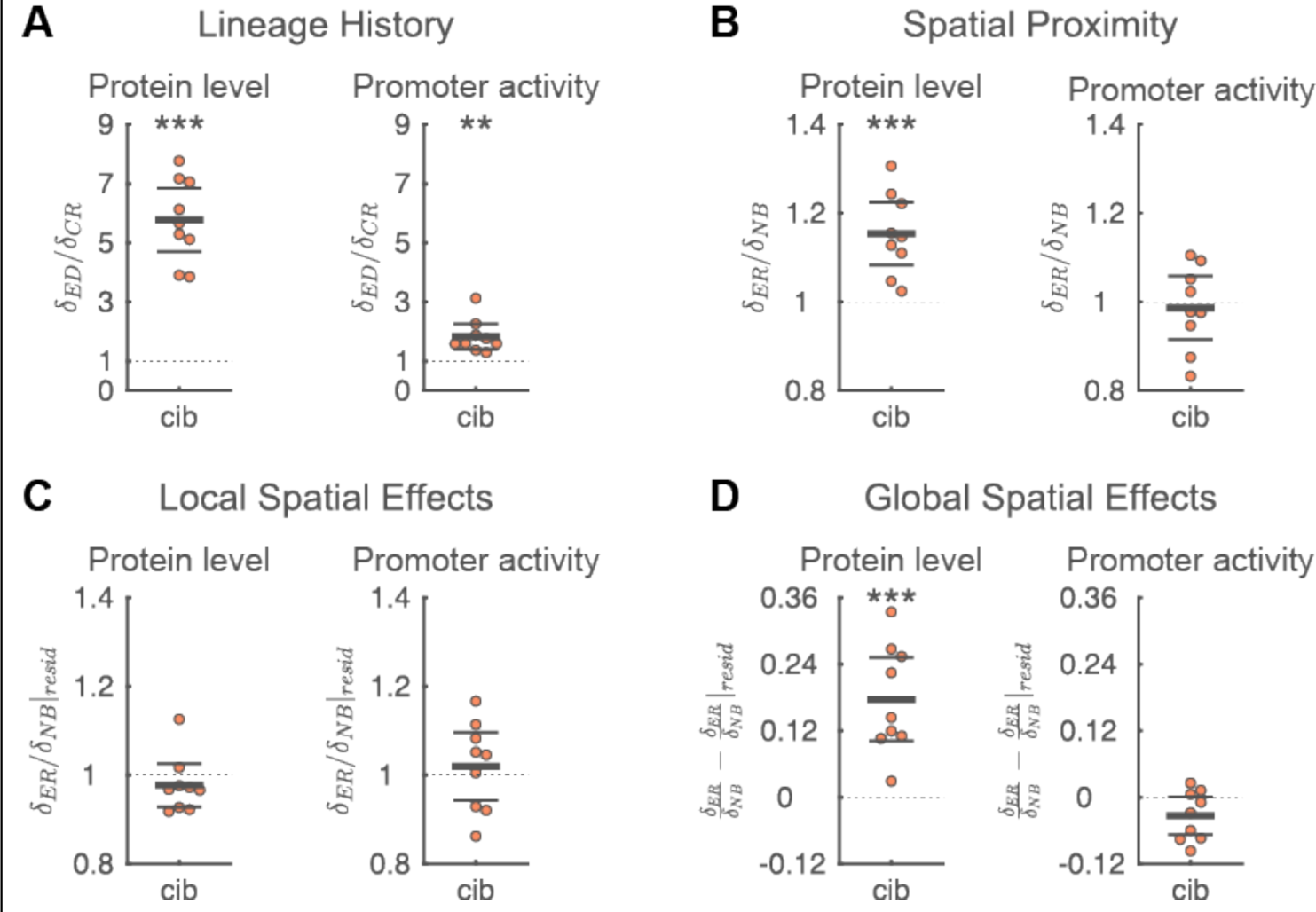
Factors contributing to spatial correlations in colicin Ib expression dynamics. **A)** Shared lineage history leads to similarity in Colicin Ib protein levels (left) and promoter activity (right). The phenotypic difference between a focal cell and an *equidistant cell* (*δ_ED_*) is significantly larger than the phenotypic difference between a focal cell and its closest relative (*δ_CR_*) i.e. 〈*δ_ED_*/*δ_CR_*〉 >1. **B)** Spatial proximity leads to similarity in Colicin Ib levels but not in promoter activity. For Colicin Ib levels, the phenotypic difference between a focal cell and an *equally-related cell* (*δ_ER_*) is significantly larger than the phenotypic difference between the focal cell and one of its neighbors (*δ_NB_*) i.e. 〈*δ_ER_*/*δ_NB_*〉 >1. **c)** Local spatial effects do not contribute to spatial correlations in Colicin Ib levels or promoter activity. Local spatial effects were calculated using the residuals of a linear regression of a cell’s phenotype to the distance of a cell to the colony edge. The difference in residuals between a focal cell and an *equally-related cell* (*δ_ER|resid_*) is not significantly different from the difference in residuals between the focal cell and one of its neighbors (*δ_ER_*/*δ_NB|resid_*), i.e. *δ_NB|resid_* ≈ 1. **D)** Global spatial effects contribute to spatial correlations in Colicin Ib levels. Global spatial effects were calculated as the difference between the total effects of spatial proximity (panel B) and the local spatial effects (panel C). **A-D)** Each point corresponds to a microcolony with 117-138 (mean=128) cells; points are horizontally offset. Thick horizontal lines indicate mean, thin lines 95% confidence intervals. Dashed lines indicate the expected value under the null-hypothesis (1 for panel A-C, 0 for panel D). Null hypothesis rejected with: ^*^p<0.05, ^**^p<0.01, ^***^p<0.001, t-test, *n*=9. The statistics are robust to the choice of the *equally-related cell* (Figure 3–Figure Supplement 1), the size of the colony being analyzed (Figure 3–Figure Supplement 2), and differences in the processing of fluorescent images (Figure 3–Figure Supplement 3). Full data can be found in Figure 3 – Source Data 1.

### Spatial proximity leads to spatial correlations in Colicin Ib protein level

Can the observed spatial correlation in Colicin Ib protein levels be fully explained by the shared lineage history of neighbors, or are there additional factors that couple gene expression in cells depending on their spatial proximity? To answer this, we investigated if neighboring cells are more similar to each other than could be expected based on how related they are. If spatial proximity by itself also contributes to spatial correlations in phenotypes, this would suggest that a cell’s phenotype is partly determined by its population context. Population context could matter both due to global spatial effects, that is, due to systematic differences in gene expression levels with a cell’s location in the colony, or due to local interactions between neighboring cells. We first analyzed whether spatial proximity by itself contributes to spatial correlations in phenotype. Subsequently, we investigated whether the contribution of spatial proximity is due to global and/or local effects.

We developed a test to quantify whether spatial proximity contributes to spatial correlations in phenotypes, after correcting for the effects of shared lineage history. We corrected for the effects of shared lineage history by comparing a group of cells that are identical in their relatedness, but that differ in how far they are apart. We then asked whether a cell is more similar to its neighbors than to cells that have the same relatedness, but that are further away in space. Specifically, we compared three cells: a focal cell, one of its neighbors, and an *equally-related cell*, i.e. a cell that has the same relatedness to the focal cell as the neighbor but that is further away in space (Figure 2C). We then compared the phenotypic difference between the focal cell and the *equally-related cell* (δ_ER_) with the phenotypic difference between the focal cell and its neighbor (δ_NB_). The effect of spatial proximity was quantified using the ratio of these two quantities (i.e. δ_ER_/δ_NB_). Values larger than 1 indicate that spatial proximity leads to similarity in phenotype (i.e. positive correlations), while values smaller than 1 indicate that spatial proximity leads to dissimilarity in phenotype (i.e. negative correlations).

We found that spatial proximity leads to significant similarity in Colicin Ib levels: the phenotypic difference between a focal cell and an *equally-related cell* is on average 1.15 times higher than the phenotypic difference between the focal cell and its neighbor (Figure 3B, 〈*δ_ER_*/*δ_NB_*〉=1.15, p=1·10^−3^. However, spatial proximity does not significantly affect promoter activity (Figure 3B, 〈*δ_ER_*/*δ_NB_*〉=0.99, p=0.7).

Why did we observe that spatial proximity leads to similarity in protein levels, but not in promoter activities? One possible reason is that protein levels have a longer autocorrelation time than promoter activities, i.e. they are influenced by events further in a cell’s past. The amount of proteins inside a cell is the sum of protein production and protein inheritance. The protein level is therefore influenced by the transcriptional activity of both the cell itself and its ancestors. Contrarily, we determined the promoter activity as the amount of proteins produced during a time scale corresponding to roughly one cell cycle. It thus reflects only the transcriptional activity within this time period. The overall (i.e. time-averaged) activities of cells could be spatially correlated even if they are not synchronized in time, for example due to transcriptional bursts or time-delays between the responses in neighboring cells. This would lead to an observed similarity over longer timescales (i.e. for protein levels) even when there is no similarity on shorter time scales (i.e. for promoter activity). Furthermore, promoter activities, which are calculated using temporal derivatives, are likely more affected by measurement noise than protein levels, which are directly measured. Weak effects of spatial proximity on similarity could thus be harder to detect using promoter activities than using protein levels.

In summary, our data shows that both shared lineage history and spatial proximity contribute to positive spatial correlations in Colicin Ib protein levels. Shared lineage history also contributes to positive correlations in *cib* promoter activity, but there is no evidence that spatial proximity also affects promoter activities.

### Global spatial effects lead to correlations in Colicin Ib protein levels

Spatial proximity could lead to correlations in expression levels in two ways: by global and by local spatial effects. Global spatial effects refer to situations where expression dynamics vary systemically with the overall position of a cell in the microcolony. Specifically, we investigated whether expression dynamics correlated with the distance of a cell to the edge of the colony. Additionally, local (i.e. microscale) effects could lead to correlations in expression dynamics. Such local effects are most likely the result of interactions between neighboring cells. These interactions can be either the result of direct sharing of cellular components, or be a consequence of intercellular feedbacks mediated though the local microenvironment. Both global and local effects could thus affect the phenotype of a cell through spatial variation in the environment. Although there is no fundamental difference between these two situations, we reserve global spatial effects for cases where the microenvironment varies on spatial scales that are (much) larger than the size of a cell and use local effects for cases where variation occurs on scales comparable to the size of a cell.

First we analyzed whether global spatial effects contribute to correlations in Colicin Ib expression dynamics. Visual inspection suggested that expression levels increased towards the center of the colonies. A linear regression of Colicin Ib protein levels with a cell’s distance to the colony edge confirmed this observation (mean r^2^=0.096, Figure 3–Figure Supplement 4). As neighboring cells are similar in their distance to the colony edge, the observed correlation could thus lead to similar expression levels in neighboring cell

To what extend does the observed global trend in colicin Ib expression dynamics explain the effect of spatial proximity? To find out, we recalculated the effect of spatial proximity after correcting for the global spatial effects. We can correct the phenotype of a cell for the global spatial effects by subtracting the expected phenotype determined by the linear regression from the observed phenotype. We then recalculated the effect of spatial proximity using these residuals (Figure 3C). This gives an estimate for the importance of local spatial effects. The importance of global spatial effects is then estimated as the difference between the effect of spatial proximity determined from the observed phenotype of a cell (*δ_ER_*/*δ_NB_*, Figure 3B) with the effect of spatial proximity determined from the residuals of the linear regression (*δ_ER_*/*δ_NB|resid_*, Figure 3C).

We found that the spatial correlations in Colicin Ib levels are strongly influenced by global spatial effects (Figure 3D, 〈*δ_ER_*/*δ_NB_* – *δ_ER_*/*δ_NB|resid_*〉=0.18, p=6·10^−4^. In fact, we no longer observe a significant effect of spatial proximity when we only analyzed the local spatial effects (Figure 3C, 〈*δ_ER_*/*δ_NB|resid_*〉=0.98, p=0.3). The spatial correlations in Colicin Ib levels are thus mainly the result of shared lineage history and shared overall position in the colony. Does this mean that intercellular feedbacks do not play any role in Colicin Ib expression patterns?

### Direct cell-cell interactions in SOS response

We designed an experimental system where we could directly test if intercellular feedbacks affect Colicin Ib expression dynamics. The system consists of two strains: an *inducible strain* in which we can induce the expression of a target gene and a *reporter strain* that has a reporter for the same gene, but that does not respond to the inducing signal. If intercellular feedbacks were present, reporter cells neighboring inducible cells should have higher expression levels than isolated reporter cells.

We expect that *cib* expression is mainly the result of fluctuations in SOS response activity, as we relieved transcriptional repression by Fur by chelating free iron in the medium (Nedialkova, Denzler, Koeppel, Diehl, Ring, Wille, Gerlach, and Stecher 2014b; Spriewald et al. 2015; Pennington and Rosenberg 2007). We thus hypothesized that any interactions in colicin expression dynamics would most likely be the result of intercellular feedbacks in SOS response. To test this hypothesis, we constructed a strain in which we could induce SOS response by expressing a nuclease inside the cell (Figure 4A, *Methods*). We combined this strain with a reporter strain that carried a transcriptional reporter for *recA*, which is a key component of the SOS response and has previously been used as a reporter for SOS induction levels (Friedman et al. 2005). The two strains were grown together on agar pads and we compared SOS induction levels in reporter cells that neighbored inducible cells with reporter cells without inducible neighbors. We found that SOS induction levels significantly increased when reporter cells neighbored an inducible cell (Figure 4B, mean relative induction=1.030, p=9·10^-4^, n=15). Although the increase in SOS induction levels is significant, its effect is rather small: reporter cells neighboring inducible cells have on average only 3% higher RecA protein levels than isolated reporter cells. One reason for the small effect size could be that SOS induction levels increased only in a small fraction of the inducible cells (Figure 4– Figure Supplement 1). When we average over all reporter-inducible cells pairs we thus underestimate the true effect size. Nonetheless, this data strongly suggests that there are intercellular feedbacks in SOS-response: cells appear to “sense” the SOS response level in their neighbors and respond by upregulating their own SOS response.

**Figure 4.**
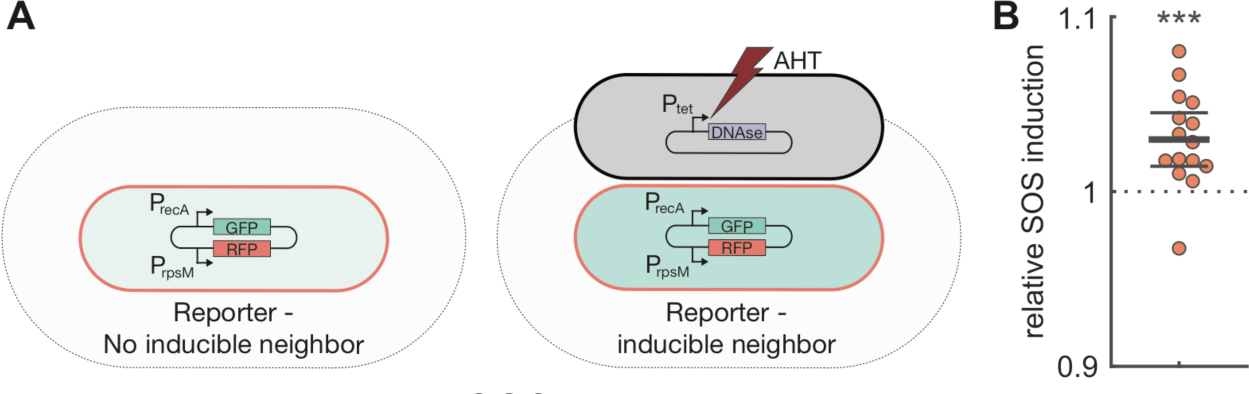
Direct cell-cell interactions in SOS response. **A)** Test for direct interactions in SOS-response. Cells with a transcriptional reporter for *recA* (pSV66-recA-rpsM, red cells) where grown together on agar pads with cells in which SOS response was induced by expressing the nuclease domain of colicin E2 (pSJB18, black cells). After 1h, the average SOS induction level was compared between reporter cells that do (right) and do-not (left) have inducible neighbors. The grey area indicates the region where cells are considered neighbors. Nuclease expression was induced by adding Anhydrotetracycline (AHT) to the agar pad. **B)**. Cells neighboring inducible cells have higher SOS response levels. Each dot shows the mean GFP intensity of a *recA* transcriptional reporter in cells with inducible neighbors divided by the mean intensity in reporter cells with no direct inducible neighbors. For each replicate we analyzed 51-189 (mean=137) reporter cells with inducible neighbors and 359-713 (mean=575) reporter cells with no inducible neighbors. Points are horizontally offset, thick horizontal line indicates mean, thin lines 95% confidence intervals. Reporter cells neighboring inducible cells have significantly higher levels of *recA* expression with a mean relative SOS induction of 1.030 (95% CI=1.015,1.045), p=9·10^-4^, t-test, *n*=15. Full data can be found in Figure 4 – Source Data 1.

Despite finding evidence for direct intercellular feedbacks in SOS response, we did not observe any effect of spatial proximity on *recA* expression (Figure 3–Figure Supplement 5). RecA protein levels are spatially correlated (Figure 1–Figure Supplement 2), however this can be fully explained by the effects of shared lineage history (Figure 3–Figure Supplement 5A 〈*δ_ED_*/*δ_CR_*〉=5.9, p=1·10^−4^). We did not find a significant effect of spatial proximity (Figure 3–Figure Supplement 5B, 〈*δ_ER_*/*δ_NB_*〉=1.07, p=0.2). One explanation for this finding is that the intercellular feedbacks are too weak to be detected with our statistical method. Another possibility is that intercellular feedbacks can only be detected when a large enough fraction of the population has high levels of SOS induction. We did not induce SOS response in the microcolonies that were used to analyze the effects of spatial proximity. The fast majority of cells in the colony had thus only very low SOS induction levels. In contrast, in our experiment where we detected evidence for interactions, we had induced high levels of SOS response in at least part of the population.

### Shared lineage history and spatial proximity lead to spatial correlations in anabolism

Next, we turn to our hypothesis that neighboring cells should be dissimilar in their expression levels of anabolic pathways. We studied three pathways involved in amino acid biosynthesis in *E. coli* and followed their expression dynamics using transcriptional reporters. Specifically, we followed: PheL, the leader peptide of the *pheLA* operon that encodes for an enzyme involved tyrosine and phenylalanine biosynthesis; MetA, an enzyme involved in methionine biosynthesis; and TrpL, the leader peptide of the *trpL*EDCBA operon that encodes for enzymes involved in tryptophan biosynthesis. Previous work has shown that *trpL* has relatively high variation in expression levels (Silander et al. 2012). We expected that higher levels of variation would facilitate cells to specialize on the production of different amino acids. Furthermore, it was previously found that cells with knockout mutations in the biosynthesis pathways of phenylalanine and methionine tend to grow well when combined with a large number of complementary knockout strains in cross feeding cultures (Mee et al. 2014; Wintermute and Silver 2010). This suggests that these two amino acids can efficiently be shared between cells, making it more likely that neighboring cells could specialize on the synthesis of different amino acids and complement each other.

First we analyzed the spatial correlations in protein levels using a randomization test, where we expected to observe negative correlations. Contrary to our hypothesis, we observed strong positive spatial correlations in the protein levels of all three pathways (Figure 1–Figure Supplement 2). An important driver of these positive correlations is the effect of shared lineage history: for all three pathways we observe that closely related cells are similar in both protein levels and promoter activities (Figure 5A, 〈*δ_ED_*/*δ_CR_*〉>1 for all pathways). However, even after correcting for lineage effects we observed that protein levels are similar between neighboring cells. For all three pathways the phenotypic difference between a focal cell and its neighbor is smaller than the phenotypic difference between the focal cell and an equally related, but more distant, cell. However, this effect is only significant for *pheL* (Figure 5B, 〈*δ_ER_*/*δ_NB_*〉>1 for all pathways). Based on protein levels we thus do not find any evidence for a division of labor in amino acid synthesis between neighboring cells.

**Figure 5.**
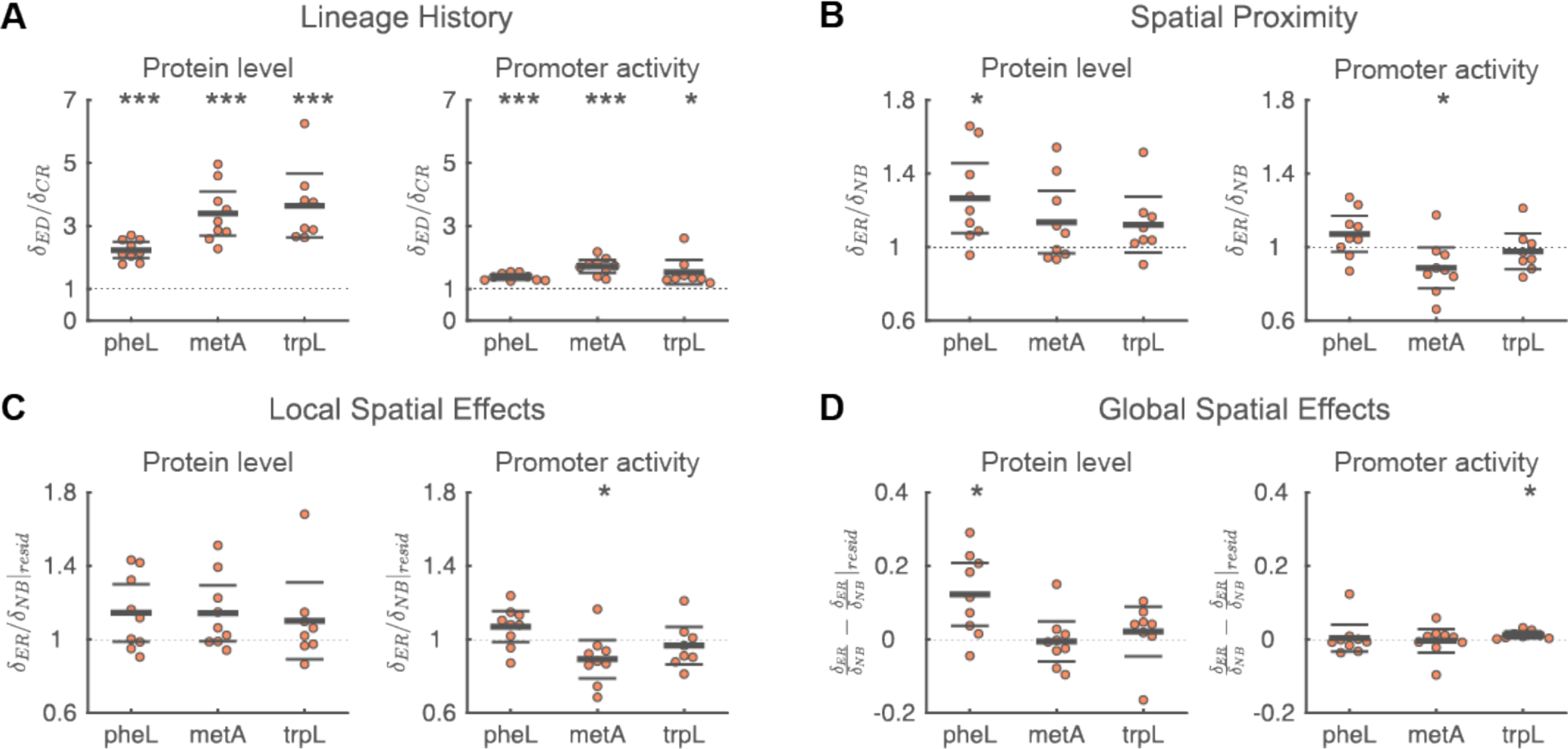
Analyses of factors that contribute to spatial correlations in amino acid synthesis. **A)** Shared lineage history leads to similarity in protein levels and promoter activity for all three pathways involved in amino acid synthesis. In all cases, the phenotypic difference between a focal cell and an *equidistant cell* (*δ_ED_*) is significantly larger than the phenotypic difference between a focal cell and itsclosest relative (*δ_CR_*) i.e. 〈*δ_ED_*/*δ_CR_*〉>1. **B)** Spatial proximity leads to similarity in PheL protein levels and dissimilarity in *metA* promoter activity. For PheL protein levels the phenotypic difference between a focal cell and an *equally-related cell* (*δ_ER_*) is significantly larger than the phenotypic difference between the focal cell and one of its neighbors (*δ_NB_*). For *metA* promoter activities, neighboring cells are less similar than expected based on their relatedness, i.e. 〈*δ_ER_*/*δ_NB_*〉<1. **C)** The dissimilarity in *metA* promoter activity is due to local spatial effects. Local spatial effects were calculated using the residuals of a linear regression of a cell’s phenotype to the distance of a cell to the colony edge. For *metA* promoter activity, the difference in residuals between a focal cell and an *equally-related cell* (*δ_ER|resid_*) is significantly smaller than the difference in residuals between the focal cell and one of its neighbors (*δ_NB|resid_*), i.e. *δ_ER_*/*δ_NB|resid_* < 1 **D)** Global spatial effects lead to similarity in PheL protein levels and *trpL* promoter activity. Global spatial effects were calculated as the difference between the total effects of spatial proximity (panel B) and the local spatial effects (panel C). **A-D)** Each point corresponds to a microcolony with 117-138 (mean=128) cells, points are horizontally offset. Thick horizontal lines indicate mean, thin lines 95% confidence intervals. Dashed lines indicate the expected value under the null hypothesis (1 for panel A-C, 0 for panel D). Null-hypothesis rejected with: ^*^p<0.05, ^**^p<0.01, ^***^p<0.001, t-test, *n*=9 (*pheL*, *metA*) or 8 (*trpL*). Full data can be found in Figure 5 – Source Data 1.

### Spatial dissimilarity in promoter activity of methionine biosynthesis

Interestingly, however, we did find negative correlations for *metA* promoter activity. This means that neighboring cells are more dissimilar in their promoter activities than expected based on how related they are. Specifically, the phenotypic difference between a focal cell and its neighbor is 1.13 times larger than the phenotypic difference between a focal cell and an *equally-related cell* (Figure 5B, 〈*δ_ER_*/*δ_NB_*〉=0.89, p=0.05). This difference does not change when we correct for global spatial effects by analyzing the residuals of a linear regression of *metA* promoter activity with the distance of a cell to the colony edge (Figure 5C, 〈*δ_ER_*/*δ_NB|resid_*〉=0.89. The dissimilarity in *metA* promoter activities is thus the result of local effects. This suggests that there are negative intercellular feedbacks affecting the expression of *metA.* In other words: if a cell transcribes *metA* at a high rate, its neighbors will tend to transcribe this gene at a low rate. For *pheA* and *trpL* we found no evidence for similar negative feedback loops in promoter activity (Figure 5B).

For *metA*, we thus observed that neighboring cells where significantly dissimilar in their promoter activities, while their protein levels tended to be similar (though the latter effect was not significant, Figure 5B). This discrepancy could be due to the different autocorrelation times of protein levels and promoter activities. At any given time, a cell could try to differentiate from its neighbors, resulting in dissimilarity in promoter activities. However, the identity and activities of a cell’s neighbors change over time due to the growth of the microcolony. The constant changes in promoter activity over time could thus prevent protein levels (which depend both on past and current promoter activities) of becoming dissimilar between neighbors.

### Spatial similarity in overall metabolic state of cells

We observed positive correlations in all three amino acid synthesis pathways we studied. This raises the question whether such positive correlations are a more general feature of a cell’s metabolism. To investigate this possibility, we simultaneously measured a cell’s elongation rate (i.e. its growth rate) and the expression level of *rpsM,* which codes for the S13 ribosomal protein. Ribosome production levels have previously been shown to be strongly correlated to a cell’s growth rate (Scott et al. 2010; Scott et al. 2014). We thus expected that *rpsM* expression is a good proxy for a cell’s overall metabolic activity.

We observed significant positive correlations in both RpsM protein levels and cell elongation rate (Figure 1–Figure Supplement 2). Overall metabolic activities thus appear to be spatially correlated. A large part of this correlation is again due to the effects of shared lineage history (Figure 6A). However, even after correcting for the effects of lineage history, we observed that neighboring cells are similar in the protein level and promoter activity of *rpsM* and in cell elongation rate (Figure 6B). Neighboring cells are thus more similar in their metabolic activities than expected based on their relatedness.

**Figure 6.**
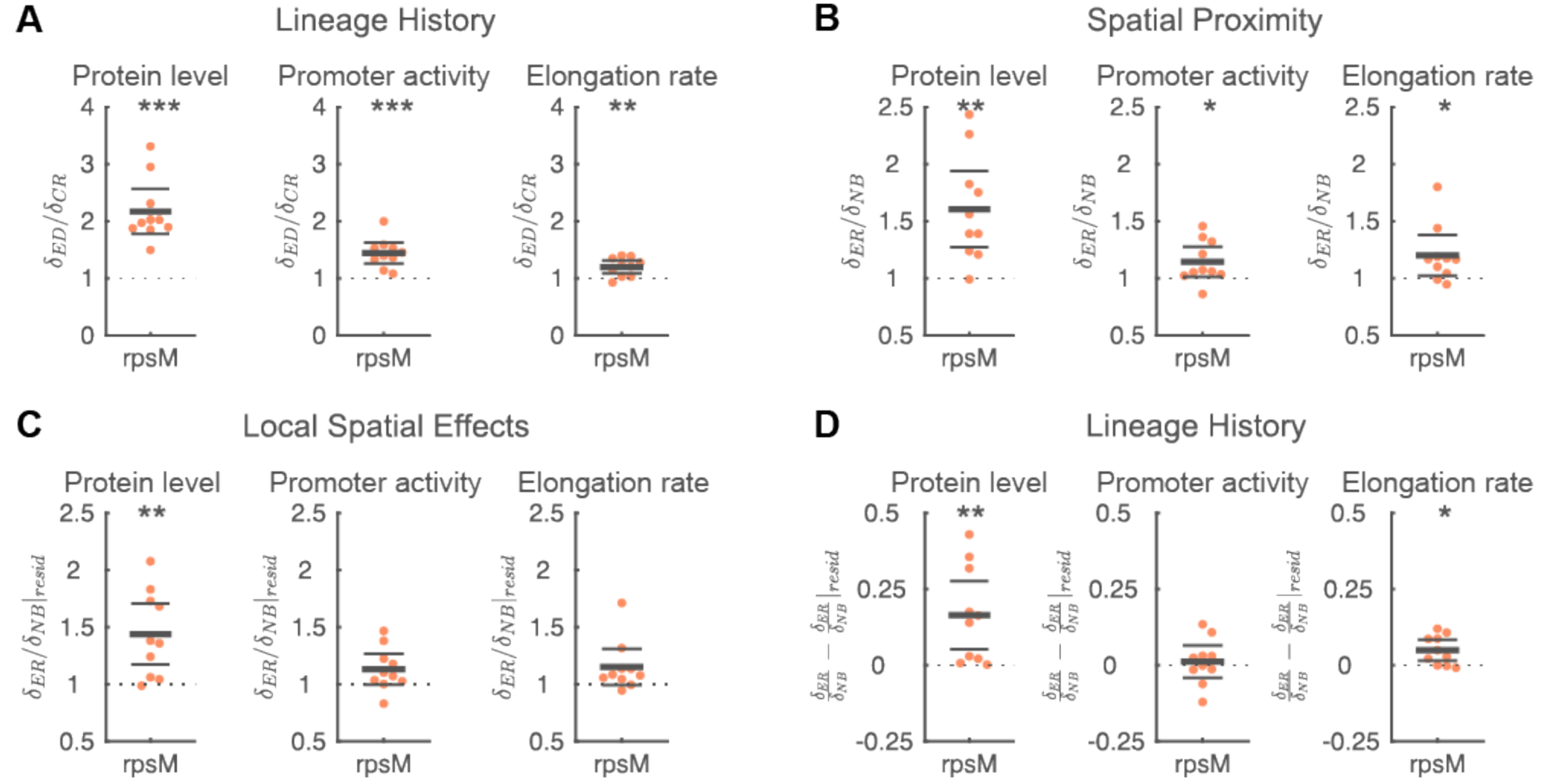
Analyses of factors that contribute to spatial correlations in metabolism. **A)** Shared lineage history leads to similarity in RpsM protein levels (left), *rpsM* promoter activity (middle), and cell elongation rate (right). In all cases, the phenotypic difference between a focal cell and an *equidistant cell* (*δ_ED_*) is significantly larger than the phenotypic difference between a focal cell and its closest relative (*δ_CR_*), i.e. 〈*δ_ED_*/*δ_CR_*〉>1. **B)** Spatial proximity leads to similarity in RpsM protein levels, *rpsM* promoter activity, and cell elongation rate. In all cases, the phenotypic difference between a focal cell and an *equally-related cell* (*δ_ER_*) significantly is larger than the phenotypic difference between the focal cell and one of its neighbors (*δ_NB_*), i.e. 〈*δ_ER_*/*δ_NB_*〉>1. **C)** The similarity in RpsM protein levels is partly due to local spatial effects. Local spatial effects were calculated using the residuals of a linear regression of a cell’s phenotype to the distance of a cell to the colony edge. For RpsM protein levels, the difference in residuals between a focal cell and an *equally-related cell* (*δ_ER|resid_*) is significantly larger than the difference in residuals between the focal cell and one of its neighbors (*δ_NB|resid_*), i.e. *δ_ER_*/*δ_NB|resid_* > 1. **D)** Global spatial effects lead to similarity in RpsM protein levels and cell elongation rate. Global spatial effects were calculated as the difference between the total effects of spatial proximity (panel B) and the local spatial effects (panel C). **A-D)** Each point corresponds to a microcolony with 117-138 (mean=128) cells, points are horizontally offset. Thick horizontal lines indicate mean, thin lines 95% confidence intervals. Dashed lines indicate the expected value under the null hypothesis (1 for panel A-C, 0 for panel D). Null hypothesis rejected with: ^*^p<0.05, ^**^p<0.01, ^***^p<0.001, t-test, *n*=10. Full data can be found in Figure 6 – Source Data 1.

Spatial proximity leads to similarity in metabolic activities due to both global and local spatial effects (Figure 6CD). RpsM protein levels and cell elongation rate are both (weakly) correlated with the distance of a cell to the edge of the colony (mean r^2^=0.07 and 0.02, respectively, Figure 3–Figure Supplement 4). Interestingly, both increase towards the center of the colony, showing that cells in the colony center have higher metabolic activities than cells at the edge. These global effects significantly contribute to the similarity between neighboring cells (Figure 6D). After removing the global effects, we still observed that spatial proximity tends to cause similarity in *rpsM* expression dynamics and cell growth rate, though the effect is only significant for RpsM protein levels (Figure 6C). Together, these data show that metabolic activities are spatially correlated; these correlations are mostly driven by shared lineage history and global spatial effects. However, we also found evidence for local spatial effects in metabolic activities. This suggests that intercellular feedbacks can couple metabolic activity and cell growth between neighboring cells.

## Discussion

We studied the expression dynamics of genes involved in a range of cellular activities in *E. coli* microcolonies and found highly non-random spatial gene expression patterns in all cases (Figure 1– Figure Supplement 1). Using a newly developed statistical method (Figure 2) we were able to show that the observed positive spatial correlations in gene expression dynamics are the result of a combination of shared lineage history and global and local spatial effects (Figure 7A).

**Figure 7.**
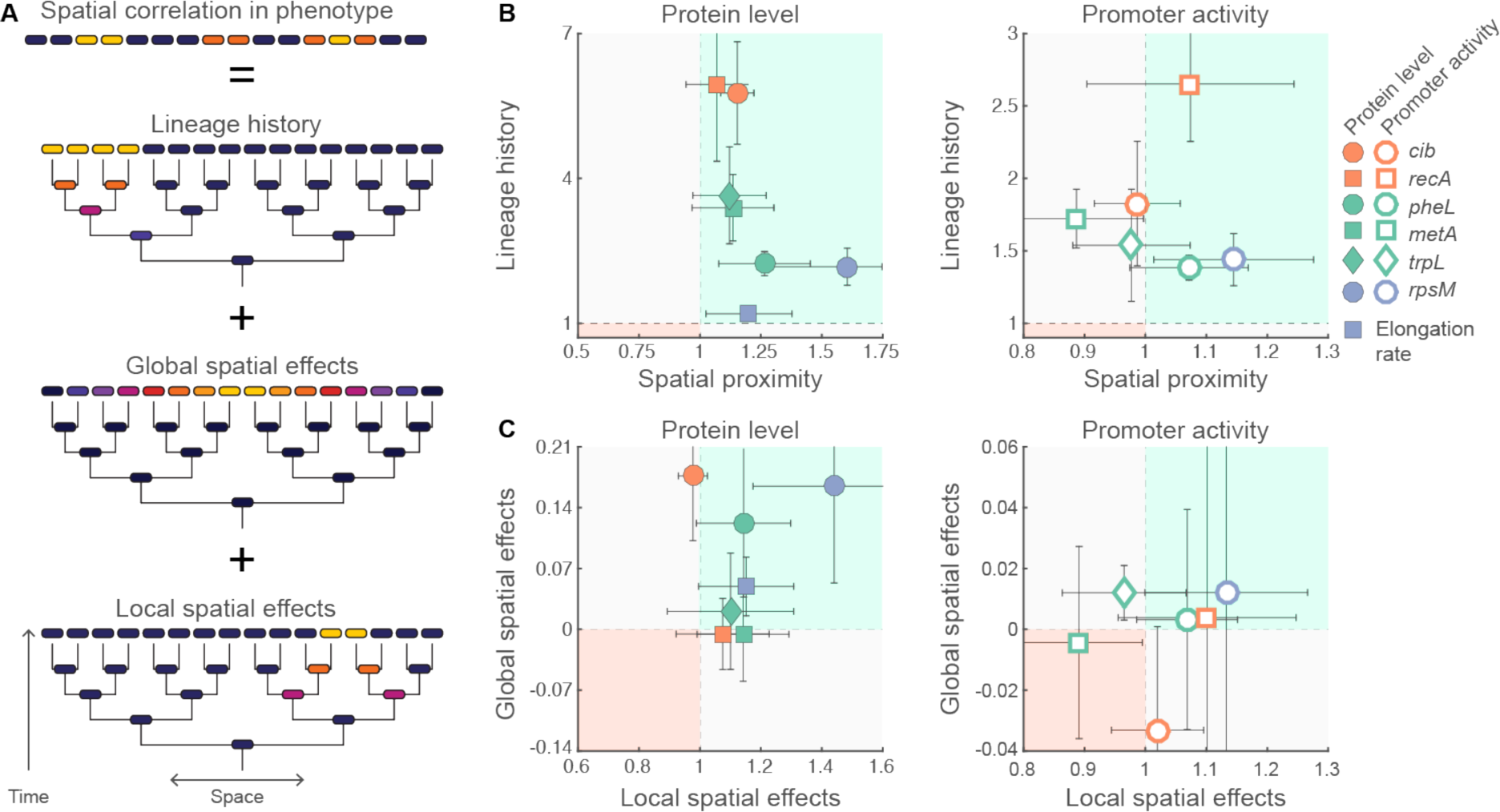
Causes of spatial correlations in phenotype. **A)** Spatial correlations in phenotype are the consequence of three major factors: shared lineage history, global spatial effects, and local spatial effects. The relative importance of these three factors differs between different pathways (see panels B and C). **B)** For each pathway the relative importance of lineage history (〈*δ_ER_*/*δ_NB_*〉) and spatial proximity (〈*δ_ED_*/*δ_CR_*〉) is shown for protein level and cell elongation rate (left) and promoter activity (right). In most cases lineage history is the dominant factor (note the different scaling of the axis). **C)** For each pathway the relative importance of global spatial effects (〈*δ_ER_*/*δ_NB_* – *δ_ER_*/*δ_NB|resid_*〉) and local spatial effects (〈*δ_ER_*/*δ_NB|resid_*〉) is shown for protein level and cell elongation rate (left) and promoter activity (right). **B, C)** Each point corresponds to the average value over 8-10 microcolonies; the data are identical to those shown in Figures 3, 5, and 6. Error bars indicate 95% confidence intervals. The green shaded region (upper right region) indicates that both factors contribute to similarity in phenotype; the red shaded region (bottom left) indicates that both factors contribute to dissimilarity in phenotype; in the other two regions (grey shading) the two factors have opposing effects.

Shared lineage history led to a similarity in protein levels and promoter activities for all studied traits (Figure 7B). Such lineage correlations have been reporter before and can easily be explained: in most cases proteins are partitioned equally between daughter cells at cell division (Robert et al. 2010; Hormoz, Desprat, and Shraiman 2015; Veening et al. 2008; Veening, Smits, and Kuipers 2008; Julou et al. 2013). With the exception of cases where cell division is asymmetric (or when proteins are present as a single copy at cell division) we thus expect shared lineage history to contribute towards positive spatial correlations.

Global spatial effects generally also contribute towards positive spatial correlations, however we found that they only affected a subset of the studied traits (Figure 7C). These global effects are likely caused by emergent spatial gradients that are the result of the uptake, release, and diffusion of metabolites during population growth (Julou et al. 2013; Stewart and Franklin 2008; Flemming et al. 2016). Interestingly, cells in the colony center grew faster and had higher expression levels of *rpsM*. This shows that nutrients are not limiting growth in the colony center, but instead that higher densities stimulate growth (possibly due to the build-up of excreted, growth promoting, metabolites). Additionally, we observed higher protein levels of Colicin Ib in the colony center. This could either be due to increased competition for iron (reducing repression by Fur) or due to higher SOS-induction levels. Although *recA* transcription levels hardly varied with the distance to the colony edge (Figure 3–Figure Supplement 4 and 5), variation in its post-translational activation could still explain the higher *cib* expression levels in the colony center.

After correcting for lineage history and global spatial effects we still observed significant effects of spatial proximity for RpsM protein levels and *metA* promoter activity (Figure 7C) and we found direct evidence for intercellular feedbacks in SOS-response (Figure 4). Together these results show that a cells phenotype can depend on that of its neighbors through intercellular feedbacks. This coupling could be the result either of a direct exchange of cellular components between cells (e.g. via nanotubes (Dubey and Ben-Yehuda 2011; Pande et al. 2015), pili (Hayes, Aoki, and Low 2010), vesicles (Schwechheimer and Kuehn 2015), or membrane fusion (Ducret et al. 2013)) or indirectly, by the conditioning of the local microenvironment. However, the molecular mechanisms behind these intercellular feedbacks are currently unknown and are likely pathway specific.

Spatial correlations in expression levels can potentially have functional consequences, for example by allowing cells to coordinate their activities with their neighbors. We hypothesized that positive correlations could be beneficial for excreted compounds, while negative correlations might be of benefit in anabolism. Consistent with the first hypothesis, we observed strong positive spatial correlations in colicin Ib expression levels; however, we did not observe negative correlations for any of the investigated amino acid synthesis pathways. Although we did observe spatial dissimilarity in *metA* promoter activity, the stronger converging effect of lineage history resulted in an overall positive correlation in expression levels. Nonetheless, our data does not completely rule out the possibility of a phenotypic division of labor in amino acid biosynthesis. The activity of many metabolic pathways is at least partly controlled by posttranslational regulation (Kochanowski, Sauer, and Noor 2015). Although transcriptional reporters can give us information about the overall activities of a cell, they cannot give any insight into the dynamic posttranslational adjustments of metabolic fluxes. For example, many amino acid synthesis pathways are regulated by end product inhibition (Chubukov et al. 2014). Negative correlations in amino acid synthesis fluxes could thus be present without corresponding differences at the transcriptional level.

Intercellular feedbacks could also be important in the context of collective information processing. Inferences about the state of the environments are fundamentally noisy. One way to increase the accuracy is by pooling measurements between a group of neighboring cells through intercellular feedbacks (Hein et al. 2015; SIMONS 2004; Berdahl et al. 2013; Popat et al. 2014). Furthermore, in cases where there is heterogeneity in either environmental cues (e.g. in case of spatial gradients) or in the sensitivity of cells to these cues (e.g. if cells differ in receptor abundances), this pooling of information could allow cells to gather better information about the environment than they can in isolation (Hein et al. 2015; Berdahl et al. 2013). We expect such processes to especially be of benefit in the context of stress response system. A cell’s survival chances might increase if it preemptively upregulates its stress response system if its neighbor is stressed, even if it is not yet exposed to any stressor itself. Consistent with this idea we found evidence for direct intercellular feedbacks in SOS response. Whether this does indeed provide a benefit, and whether such feedbacks apply more generally to stress response systems are interesting directions for future work.

The functional consequences of spatial correlations in phenotypes do not directly depend on the processes that give rise these correlations. Shared lineage history, global spatial effects, and positive intercellular feedbacks could, for example, all lead to similar spatial gene expression patterns. However, there are important differences between these processes that can affect functionality. The relative positioning of a cell and its relatives is largely determined by the physics of cell growth (Nadell et al. 2013). Likewise, global spatial effects will largely be determined by the physical and chemical properties of the environment (Stewart and Franklin 2008). Both these processes are thus beyond the control of the cell and, importantly, can result in very different spatial patterns depending on environmental conditions. In contrast, intercellular feedbacks would allow cells to coordinate activities with their neighbors irrespective of the environment. This allows for more robust and consistent pattern formation. Furthermore, intercellular feedbacks allow for more flexibility as it can give rise to both positive and negative correlations and could even operate between genetically unrelated cells. Finally, intercellular feedbacks directly link the genotype of a cell to the spatial patterns of gene-expression at the colony level, potentially allowing for these patterns to evolve by natural selection.

Our most important conclusion is that the phenotype of a cell depends to a large extend on the population context in which it grows. Our work thereby joins a growing number of recent studies showing that a large degree of phenotypic variation is not random, but rather determined by a cells lineage history, location in the population and/or the activities of its neighbors (Snijder and Pelkmans 2011; Symmons and Raj 2016). We found that population context affected a number of diverse pathways, suggesting that it could be equally important in other pathways, both in *E. coli* and in other microorganisms. Furthermore, we expect the same principles to hold in more complex 3D-biofilms. If we want to understand the activities of individual cells in such systems, or the functioning of the population as a whole, it is thus essential to learn more about the feedbacks between a cell and the surrounding population.

## Methods

### Strains and reporter plasmids

All experiments were done using *E. coli* MG1655 (see *Supplementary File 1 Table 1* for a list of strains and plasmids used in this study). Gene expression dynamics were followed using transcriptional reporters. The promoter region of the gene of interest was inserted in front of a *gfpmut2* green fluorescent protein. For *trpL*, *metA*, *pheL*, and *rpsM* we used reporters based on the pUA66/pUA139 low copy number plasmids (Zaslaver et al. 2006). For *recA* we used a newly constructed dual reporter plasmid, pSV66-recA-rpsM, which is based on the low copy number plasmid pUA139 (Zaslaver et al. 2006). This plasmid contains a GFPmut2 transcriptional reporter for *recA* and an additional turboRFP (red fluorescent protein) transcriptional reporter for *rpsM*, allowing for the independent measurement of two promoters within the same cell (see *Supplementary File 2* for construction details). For *cib* we used the medium copy number plasmid pM1437 with a pBR322 background (Nedialkova, Denzler, Koeppel, Diehl, Ring, Wille, Gerlach, and Stecher 2014a; Spriewald et al. 2015). MG1655 does not naturally contain the *cib* operon in its chromosome. To measure Colicin Ib expression dynamics, we therefore transformed TB60 (MG1655 containing a chromosomal kanR cassette) with the p2-camR plasmid that is based on the natural occurring *Salmonella* pColB9 plasmid, which contains, among others, the *cib* operon (Stecher et al. 2012). We subsequently transformed the same strain with the pM1437 plasmid containing the *cib* transcriptional reporter. Additionally, we used the high copy number plasmids pGFP and pRFP with inducible green and red fluorescent proteins under control of the *lac* promoter to check for fluorescent bleed-through and halos (see *Supplementary File 1-Table 1
*).

**Table 1.**
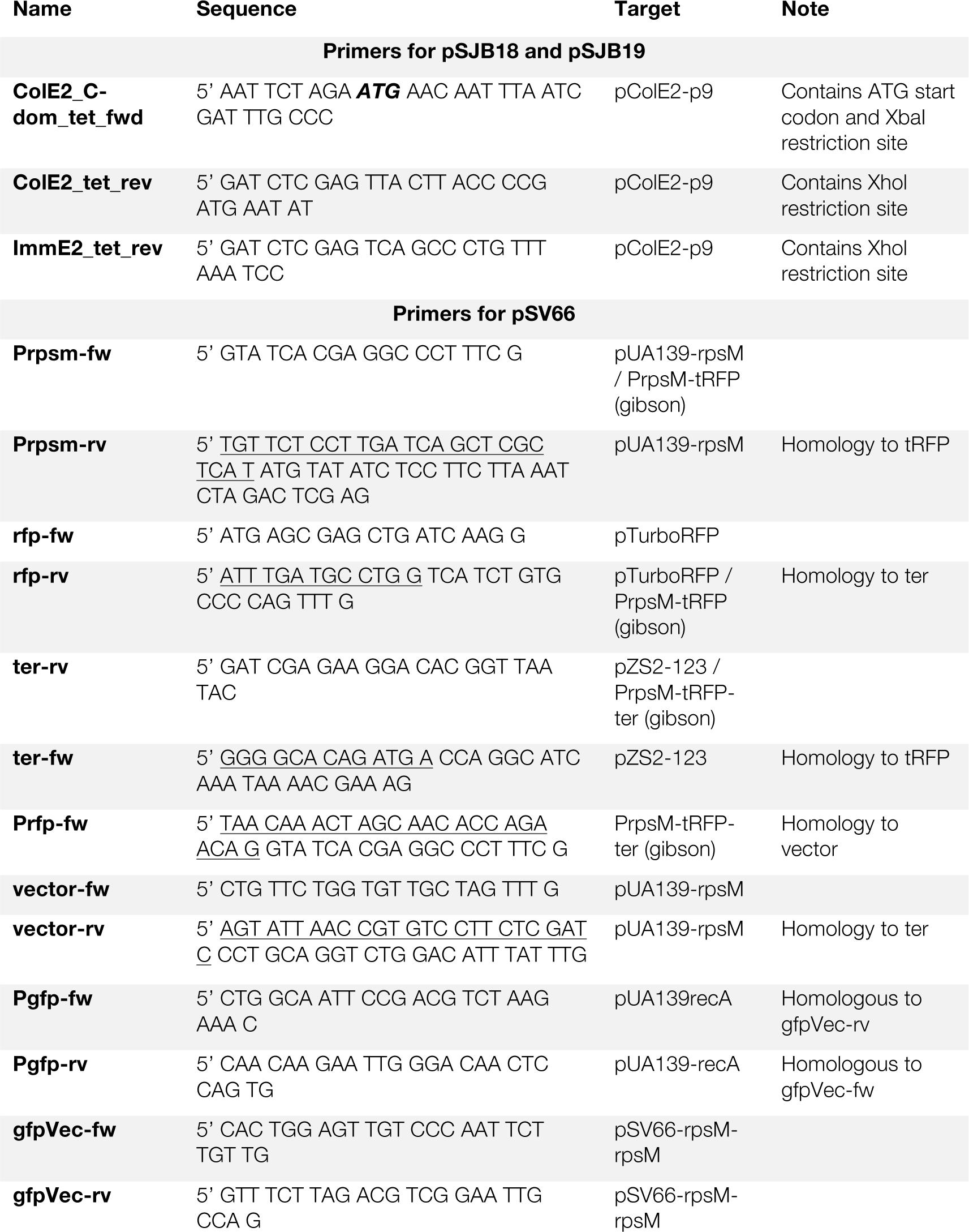
Underlined parts are tails added to primers to provide homology for Gibson assembly. Abbreviations: (gibson): primers are used to amplify end product of Gibson assembly; tRFP: turboRFP; ter: terminator

### Strain with inducible SOS response

To test for interactions in SOS response we constructed a plasmid, pSJB18, with which SOS response can be chemically induced. To do so, the nuclease domain of colicin E2 (colE2) was cloned downstream of the P_*tet*_ tetracycline inducible promoter of the pMG-Ptet vector (see *Supplementary File 2* for construction details). Upon induction, the nuclease activity results in DNA breaks, which in turn activates the cell’s SOS response. We made sure that the nuclease produced in a cell could not directly affect neighboring cells in two ways: i) pSJB18 does not contain the lysis gene that is part of the full *colE2* operon; as colicins are released during cell lysis this greatly reduces the amount of extracellular nuclease (Cascales et al. 2007). ii) pSJB18 only contains the C-terminal nuclease domain of the *colE2*; the N-terminal and central domains that are required for Colicin E2 to enter a target cells (by mediating receptor binding and membrane translocation, respectively (Cascales et al. 2007)) were removed. A second plasmid, pSJB19, was constructed. This plasmid is identical to pSJB18, except that it also contain the coding sequence for the Colicin E2 immunity protein. Expression of this immunity protein inhibits the nuclease activity of Colicin E2 (see *Supplementary File 2* for construction details).

We confirmed the functionality of the construct by co-transforming MG1655 with pSJB18 and pUA139-recA (Zaslaver et al. 2006). The latter contains a GFP transcriptional reporter for the SOS-response gene *recA*. Expression of the nuclease was induced by adding 100ng/ml of the non-toxic tetracycline analog anhydrotetracycline (AHT, Fluka, Buchs, Switzerland). Using flow cytometry and single-cell microscopy we confirmed that SOS-response activity (measured as *recA* expression levels) increased when the inducer was added (Figure 4–Figure Supplement 1).

### Media and growth conditions

In all cases cultures were started from a single colony taken from a LB-agar plate and grown overnight at 37°C in a shaker incubator. Subsequently the cultures were diluted 100 to 1000 fold and grown until mid-exponential phase. Cells containing reporters for *cib*, *recA*, and *rpsM* were grown in LB media (Sigma-Aldrich, Buchs, Switzerland, or Applichem, Darmstadt, Germany). For these reporters, microscopy was done on agar pads consisting of LB with 1.5% agar (Sigma-Aldrich or Applichem). Cells containing reporters for *metA*, *pheL* and *trpL* were grown overnight in M9 medium (47.76 mM Na_2_HPO_4_, 22.04 mM KH_2_PO_4_, 8.56 mM NaCl and 18.69 mM NH _4_Cl) supplemented with 1mM MgSO_4_, 0.1 mM CaCl_2_, 0.4% Glucose (all from Sigma-Aldrich), and 5% LB. Diluted cultures were grown in M9 medium supplemented with 1mM MgSO_4_, 0.1 mM CaCl_2_ and 0.4% Glucose. Microscopy was done on agar pads consisting of M9 salts with 1.5% agar and supplemented with 1mM MgSO_4_, 0.1 mM CaCl_2_ and 0.4% Glucose.

Plasmid maintenance was insured by adding the appropriate antibiotic to the culture medium and agar pads: 50μg/ml ampicillin (pM1437, pSJB18, pGFP, pRFP, Applichem), 50μg/ml kanamycin (pUA66, pUA139, pSV66, Sigma-Aldrich) and 15μg/ml chloramphenicol (p2-camR, Sigma-Aldrich). For experiments with the cib reporter we added additionally 0.1mM DTPA (diethylenetriaminepentaacetic acid, Fluka) to the medium of the diluted cultures and to the agar pads to chelate free iron. For the SOS interaction experiments 100ng/ml of anhydrotetracycline (AHT) was added to the agar pads to induce the nuclease in pSJB18. For these experiments, no antibiotics were added to the agar pads as the two strains (MG1655+pSJB18 and MG1655+pSV66-recA-rpsM) carry different resistance genes. As these experiments only lasted 1h, we expect plasmid loss to be negligible. For experiment using pGFP or pRFP we added 1mM IPTG (Isopropyl β-D-1-thiogalactopyranoside, Promega, Madison, Wisconsin) to the liquid cultures and agar pads to induce expression of the fluorescent proteins.

### Agar pad preparation

Agar pads were prepared by adding the appropriate supplements to molten aliquots of LB or M9 agar and adding 250µl of this mixture to the well of hollow-well microscope slides (Karl Hecht GmbH, Sondheim, Germany). The wells were sealed with a cover glass and dried at room temperature for 20 to 30min. Subsequently the cover glass was removed and the agar was cut into a square of approximately 5x5mm in the center of the well. 0.5 to 2µl of prepared cell suspension (see below) was added to the center of the pad and left to dry. Finally, the pad was sealed by adding a new cover glass. An air-tight seal was insured by adding a thin layer of lubricating grease (Glisseal, Borer, Zuchwil, Switzerland) between the two glass surfaces. The agar pad only occupies the central part of the well; the remaining area contains air to insure that sufficient oxygen is present for aerobic growth.

Before inoculation the optical density at 600nm (OD600) of the cultures was measured. The cultures were diluted to the desired OD600 (of 0.001 to 0.01) before adding 0.5 to 2µl of cells to the pad. For the SOS interaction experiment the two strains were first washed to remove antibiotics from the growth medium. Subsequently the strains were mixed in approximately a 1:1 ratio and added to the agar pad.

### Microscopy

Time-lapse microscopy was done using fully-automated Olympus IX81 inverted microscopes (Olympus, Tokyo, Japan). Imaging was done using a 100X NA1.3 oil objective (Olympus) and either a F-View II CCD camera (for *cib*, Olympus Soft Imaging Solutions, Münster, Germany) or an ORCA-flash 4.0 v2 sCMOS camera (all other data, Hamamatsu, Hamamatsu, Japan). Fluorescent imaging was done using a X-Cite120 120 Watt high pressure metal halide arc lamp (Lumen Dynamics, Mississauga, Canada) and Chroma 49000 series fluorescent filter sets (N49002 for GFP and N49008 for RFP, Chroma, Bellows Falls, Vermont). Focus was maintained using the Olympus Z-drift compensation system and the entire setup was controlled with either the Olympus CellM or CellSens software. The sample was maintained at 37^°^C by a microscope incubator (Life imaging services, Basel, Switzerland). Images were taken every 3 (*rpsM*, elongation rate), 5 (*cib*) or 7.5 (recA, *trpL*, *pheA*, *metA*) minutes for several hours.

### Selection and analysis of microcolonies

Fiji (Schindelin et al. 2012) was used for data visualization, image cropping and file-type conversions. All other processing was done using Matlab (version 2013 and newer, MathWorks, Natick, Massachusetts). For each reporter we analyzed 8 to 10 microcolonies, we decided on this sample size based on preliminary experiments with a colicin Ib reporter strain. Each micro colony is considered to be an independent biological replicate. From each agar pad we selected 1 to 6 (median=2) positions that contained an isolated micro colony of sufficient size (>128 cells before cell overlap occurs) and good optical quality. We manually determined the time range where cells were present in a single layer and cropped the images to only contain the area occupied by the colony.

Subsequently Schnitzcells 1.1 (Young et al. 2011) was used to segment and track cells. For *cib*, segmentation was done on phase contrast images. For *trpL*, *metA*, *pheA*, and *rpsM* segmentation was done on the GFP fluorescent images. The reporter plasmid for *recA* also contained a RFP reporter for *rpsM*, here segmentation was done on the RFP fluorescent images. As a last step custom Matlab code was used to extract fluorescent and geometrical properties of each cell (see fluorescent image processing and cell length determination below).

### Fluorescent image processing

There are a number of optical artifacts that could cause neighboring cells to have similar fluorescent intensity levels, it is thus essential to correct for these artifacts before calculating the spatial similarity. Specifically, we applied the following corrections (see also Figure 1–Figure Supplement 1):

- Shading correction. We corrected for inhomogeneities in the illumination field using a shading image. This image gives the normalized intensity of the incoming light for each pixel in the image. Subsequently we obtained the shading corrected image by dividing the intensity in each pixel of the captured fluorescent image by the intensity in the corresponding pixel of the shading image.
- Deconvolution. Diffraction will cause the light of a point source to be spread across several pixels. Bright cells will thus generate a “halo” that increases the fluorescent intensity of its neighbors. We corrected for diffraction by deconvolving the shading corrected image with the experimentally measured point spread function (PSF) of the microscope (Kiviet et al. 2014). Deconvolution was done using the Matlab function “deconvlucy”, which uses the Lucy-Richardson method. We found that the accuracy of the deconvolution correction depends critically on the size of the PSF that is used (Figure 1–Figure Supplement 3B). When the size of the PSF is too small (e.g. 13×13 pixels) the halos are not completely removed, increasing fluorescent intensities in neighbors of bright cells. If the PSF is too large (e.g. 30×30 pixels) a “dark halo” artifact is formed, decreasing fluorescent intensities in neighbors of bright cells. We calibrated the required size of the PSF by mixing unlabeled wild type cells with cells carrying a high copy number plasmid with an inducible green fluorescent protein. We then selected the size of the PSF (i.e. 24×24 pixels) for which median fluorescent intensity in unlabeled cells neighboring GFP labeled cells are the same as for isolated unlabeled cells (Figure 1– Figure Supplement 3B,C). Furthermore, we confirmed that our statistical analysis of the effects of spatial proximity is robust to small changes in the size of the PSF (Figure 3–Figure Supplement 3).
- Background correction. We performed a background correction to compensate for temporal changes in the incoming light intensity. For each pixel the background corrected intensity (*I*_*corr*_) was calculated as: *I*_*corr*_ = (*I-Dark*)/(*Bg*-*Dark*), where *I* is the pixel intensity after shading correction and deconvolution, *Bg* is the median intensity of all background pixels (i.e. all pixels that are not part of any segmented cell) and *Dark* is the median pixel intensity for the dark image (i.e. an image taken when no light reaches the camera taken with the same exposure settings).
- Cell center intensity. As sell segmentation is imperfect, some pixels at the periphery will be misclassified. To increase robustness to such errors we calculated the mean fluorescent intensity only over the central area of the cell. The central area is found by eroding (i.e. shrinking) the cell segmentation mask on all sides with one quarter of the median cell width (the median cell width was determined over all cells in the microcolony). For most cells the intensity was thus be determined for the central 50%. If erosion removed all pixels in the cell mask, we progressively reduced the number of outer pixels we removed until at least a single row of pixels remained in the cell center.
- Fluorescence bleed-through. The *recA* reporter strain contained a second RFP reporter for *rpsM*. We confirmed that there is no fluorescence bleed-through from the RFP to the GFP channel. To do so, we mixed unlabeled cells with cells carrying a high copy number plasmid with inducible red fluorescent protein. The distribution of fluorescent intensities in the GFP channel in unlabeled cells is identical to intensities in the GFP channel for the RFP labeled cells. This shows that emission from the red fluorescent protein do not affect measured intensities in the GFP channel (Figure 1–Figure Supplement 3A).

After performing all corrections, we obtained for each cell the mean fluorescent intensity, *Ī*(*t*) which is proportional to the concentration of GFP molecules in the cell and hence to the concentration of the gene of interest. Throughout the text we use protein level to refer to the mean fluorescent intensity.

### Cell length determination

Cell length was determined following the procedure described in (Kiviet et al. 2014). In short: the cell centerline was determined by fitting the cells mask with a 3th degree polynomial (*f*(*x*)) To find the cell pole positions we calculated the silhouette proximity (sum of the squared distance to closest 25 pixels in cell mask) along the centerline. This measure is constant in the cell center, but increases sharply at the poles; the position of the cell poles was taken as the point along the centerline where the proximity silhouette reached 110% of the average value in the cell center. The cell length was subsequently calculated by numerical integration of 
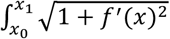
 where *f'*(*x*) is the derivative of *f*(*x*) and *x*_0_ and *x*_1_ are the positions of the cell pole.

### Cell elongation rate

Cell elongation rates (*r*) were calculated for the microcolonies with a *rpsM* transcriptional reporter by fitting the exponential curve *L*(*t*) = *L*(0) · *e^r·t^* to the cell length over time. The fitting was done using a linear fit on the log transformed cell lengths over a sliding time window of 7 time-points (21 minutes). When the time window exceeded the life time of a cell, it was extended by summing the cell lengths of the two daughter cells or by taking a fraction of *L*_0_/(*L_0_* + *L*_0,*sister*_) of the mother cell length. Here, *L*_0_ and *L*_0,*sister*_ are the lengths of a cell and its sister at their birth. This fraction takes the effects of asymmetries at cell division into account.

### Promoter activity

The promoter activity (PA) is estimated as the rate of change in the total fluorescent intensity of a cell: 
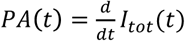
 The measurement of total fluorescent intensity as the summed intensity over all pixels in the cell mask is very sensitive to segmentation inaccuracies. To get a more accurate estimate of promoter activities we thus estimate the total fluorescent intensity by multiplying the mean fluorescent intensity of a cell, *Ī*(*t*) with its length, *L*(*t*); as the cell width is constant through the lifetime of a cell, this quantity is proportional to the total fluorescent intensity of a cell: *I_tot_*(*t*) ∝ *Ī*(*t*) · *L*(*t*) = *Ī*(*t*). The promoter activity is then estimated as the slope of a linear fit of this quantity over a window of 5 time points: 
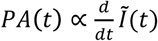
. When the time window exceeded the life time of a cell, it was extended by summing the total fluorescent intensities of the two daughter cells or by taking a fraction 
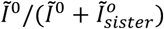
 of the total intensity of the mother cell. Here, *Ī*^0^ and 
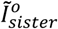
 are the total fluorescent intensities of a cell and its sister at their birth. This fraction takes the effects of asymmetries at cell division into account.

### Cell geometric and neighborhood properties

The cell width and its position were determined using the Matlab *regionprops* function and correspond to the minor-axis length of a fitted ellipse and the coordinates of the center of mass, respectively. The neighbors of a cell were found by expanding the cell mask in all directions with ¾ of the median cell width; all cells that overlap with this expanded area are classified as neighbors. The distance of a cell to the colony edge was determined as the minimum Euclidean distance between pixels inside the cell mask and pixels that are part of the colony boundary.

### Neighborhood similarity statistic

We quantitatively investigated the apparent non-randomness of expression patterns in the microcolonies using a randomization procedure. Cells in the colony were classified into two groups depending on whether their mean fluorescent intensity was above or below the median intensity in the microcolony. For each cell in the colony we calculated the fraction of neighboring cells that was classified in the same group and we computed the mean over all cells (red bar, Figure 1D, – Figure Supplement 2). We then randomly permuted intensities between cells in the colony and recalculated the mean fraction of neighbors classified in the same group. This procedure was repeated 10^4^ times obtaining the distribution shown in Figure 1D. p-values were calculated by taking the fraction of randomized samples that have a higher mean fraction of neighbors of the same type than the non-randomized data.

### Statistic for effect of shared lineage history

We tested for the effect of shared lineage history by quantifying how similar a cell is to its closest relative after correcting for the effects of spatial proximity. We compared the phenotypes within a group of three cells: a focal cell, its closest relative, and an *equidistant cell*. The closest relative will typically be a cell’s sister, however if the sister has already divided we selected one of its offspring (e.g. a niece of the focal cell) at random. The *equidistant cell* is a cell that directly neighbors the closest relative and that has a center-to-center distance to the focal cell that is the most similar to that of the closest relative.

We then calculated the difference in phenotype between the focal cell *i* and its closest relative (*CR_i_*): 
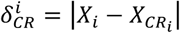
 and between the focal cell *i* and its *equidistant cell* 
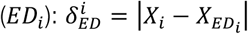
 where *X_i_*, *X_CR_i__*, and *X_ED_i__* are the phenotypes (i.e. protein levels, promoter activities, or elongation rates) of the focal cell, closest relative, and *equidistant cell*, respectively. Finally, we calculated the effect of shared lineage history by taking the ratio of these two phenotypic distances: 
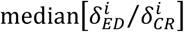
 where the median is taken over all cells in the colony.

### Statistic for the effect of spatial proximity

We tested for the effect of spatial proximity by quantifying how similar a cell is to its neighbors after correcting for the effects of shared lineage history. We compared the phenotypes within a group of three cells: a focal cell, one of its neighbors, and an *equally-related cell*. We defined a cell’s neighbors as all cells that are directly adjacent (within 3/4 cell width) to the focal cell (mean number of neighbors=5,95% range=[3,8]). For each neighbor we found a group of *equally-related cells*, these are cells that have the same relatedness to the focal cell as the neighbor, but that are further away in space (mean number of *equally-related cells*=20,95% range=[0,70]). From this group we selected the most distant *equally-related cell*, which is the *equally-related cell* with the largest Euclidean distance to the focal cell.

We then calculated the difference in phenotype between the focal cell *i* and its neighbor *j*: 
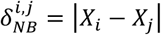
and between the focal cell and the most distant *equally-related cell*

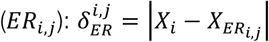
 where *X_i_*, *X_j_*, and *X_ER_i,j__* are the phenotypes (i.e. protein levels, promoter activities, or elongation rates) of focal cell *i* its neighbor *j*, and their most distant *equally-related cell*. Finally, we calculated the effect of spatial proximity by taking the ratio of these two phenotypic distances: 
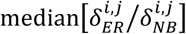
 where the median is taken over all neighbor-focal cell pairs in the colony.

We also tested whether our choice of *equally-related cell* affected our conclusions (Figure 3–Figure Supplement 1). We recalculated the statistics using the median phenotypic distance between the focal cell and all *equally-related cells*: 
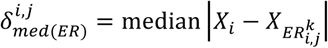
 where 
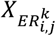
 is the phenotype of the *k*^th^ *equally-related cell* of neighbor *j* of focal cell *i* and the median is taken over all *equally-related cells*. Subsequently we calculated the effect of spatial proximity as: 
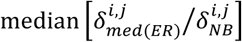
 where the median is taken over all neighbor-focal cell pairs in the colony.

### Statistic for local spatial effects

The phenotype of a cell can vary systematically within the colony, we corrected for such global effects using a linear regression of a cell’s phenotype (*x*_*i*_) with its distance to the edge of the colony (*d*_*i*_): *X*_i_ = *α* + *β* · *d*_i_ where *α* and *β* are constants. The strength of local spatial effects could then be estimated by correcting the the observed phenotype of a cell (
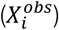
) for the global trend by calculating the residuals of the regression: regression: 
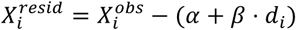
. We then calculated the ratio of phenotypic differences 
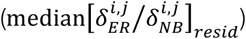
 as described above, where the phenotype of a cell (*X*_*i*_) was replaced with the residual of the regression 
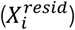

### Statistic for global spatial effects

The importance of global spatial effects was quantified by calculating to what extend the effect of spatial proximity is reduced when we correct for the systematic variation in phenotype. Specifically, we defined the global spatial effect as: 
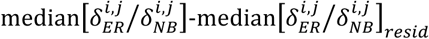
 whµere the first term describes the total effect of spatial proximity and the second term describes the local spatial effects.

## Acknowledgements

We thank Daan Kiviet, Alejandra Manjarrez, Alejandra Rodriguez, Ben Roller, Susan Schlegel, and Clément Vulin for helpful comments and discussion on a previous version of this manuscript and Daan Kiviet and Roland Mathis for providing parts of the Matlab code used in this work.

### Competing interests

The authors declare no competing interests.

## Figure Supplements

**Figure S1.**
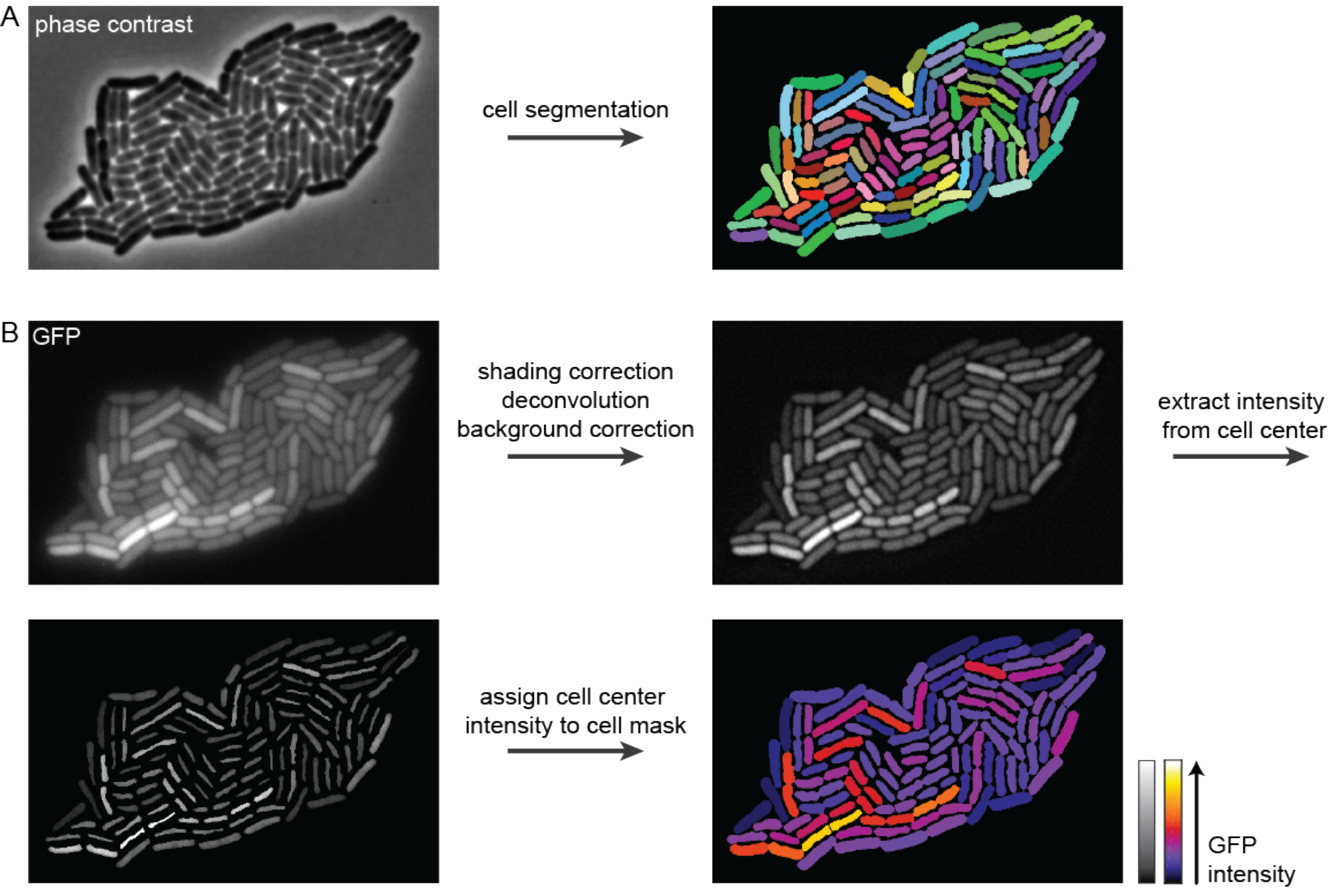
Figure 1–Figure Supplement 1. Image processing pipeline. **A)** Phase contrast images were segmented using Schnitzcells (Young et al. 2011) to find the cell masks (right). For some reporters, segmentation was done on fluorescent images (see *Methods*), **B)** GFP images (upper left) were corrected for inhomogeneities in the illumination field (shading 7 correction), diffraction (deconvolution), and background fluorescence (background correction), the resulting corrected image is shown (upper right). The mean fluorescence intensity was determined in the cell center only (lower left) and assigned to the entire cell mask (lower right). The figure shows an 10 example of a microcolony with a transcriptional reporter for *cib*

**Figure S2.**
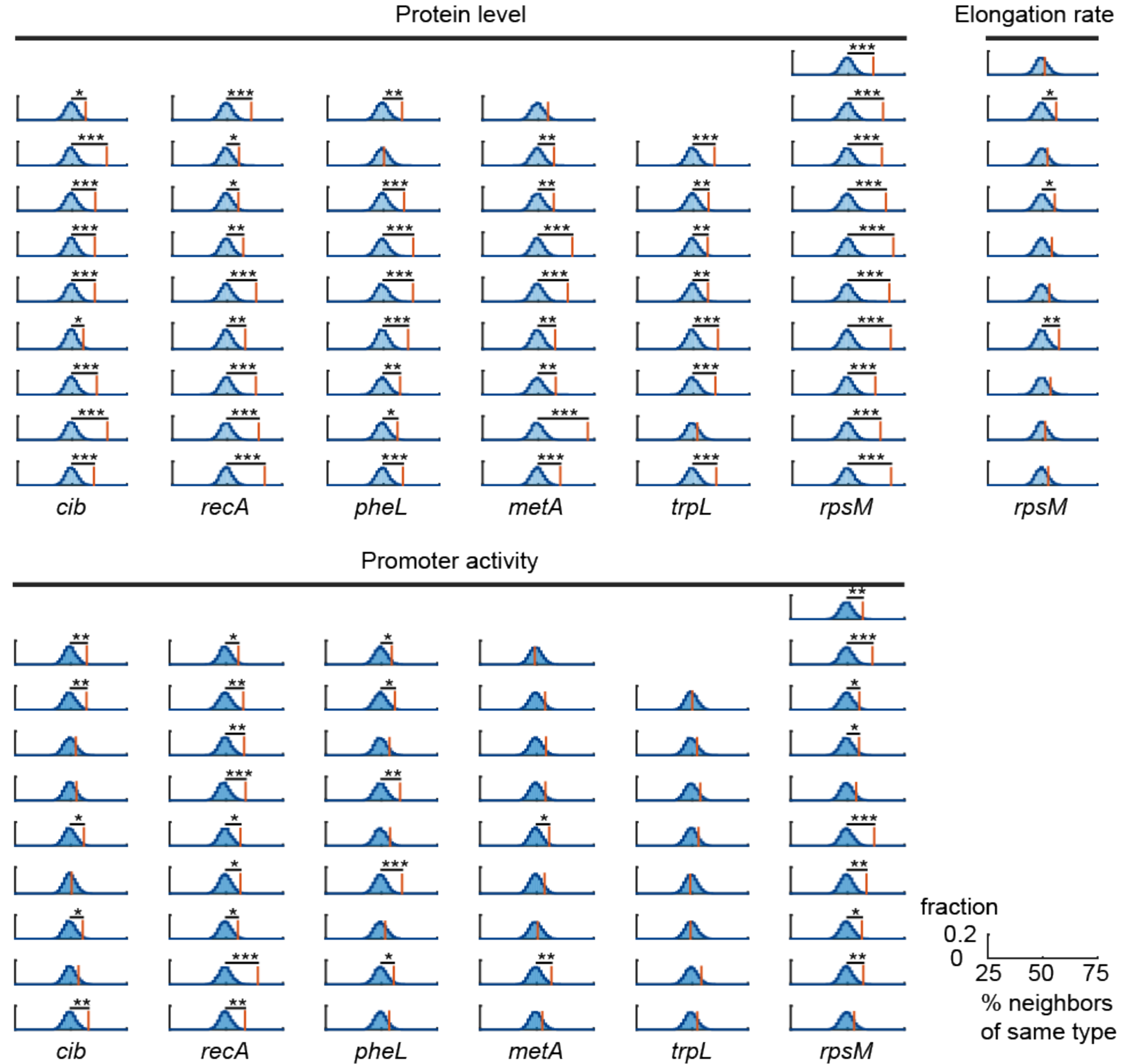
Figure 1–Figure Supplement 2. Neighboring cells have similar expression levels. For all measured phenotypes (except *trpL* promoter activity) we observed that neighbors are significantly more similar than can be expected by chance (p<10^-6^, except for: elongation rate (p=3·10^-5^), *metA* promoter activity (p=3·10^-4^), and *trpL* promoter activity (p=0.05), Chi-squared test on combined p-Values using Fischer’s method). Spatial correlations in phenotypes were quantified using a randomization test: cells were grouped into two clusters (intensity above or below median intensity). Each cell was compared to all its neighbors and the percentage of neighbors of the same cell type was calculated (red line). This procedure was repeated 10^4^ times after randomly permuting measured intensities among all cells (blue distribution). Each column shows 8 to 10 replicate microcolonies of 117-138 (mean=128) cells for each reporter. Data is shown for three phenotypes: protein level (upper left), elongation rate (upper right), and promoter activity (bottom). *p<0.05, **p<0.01, ***p<0.001, randomization test.

**Figure S3.**
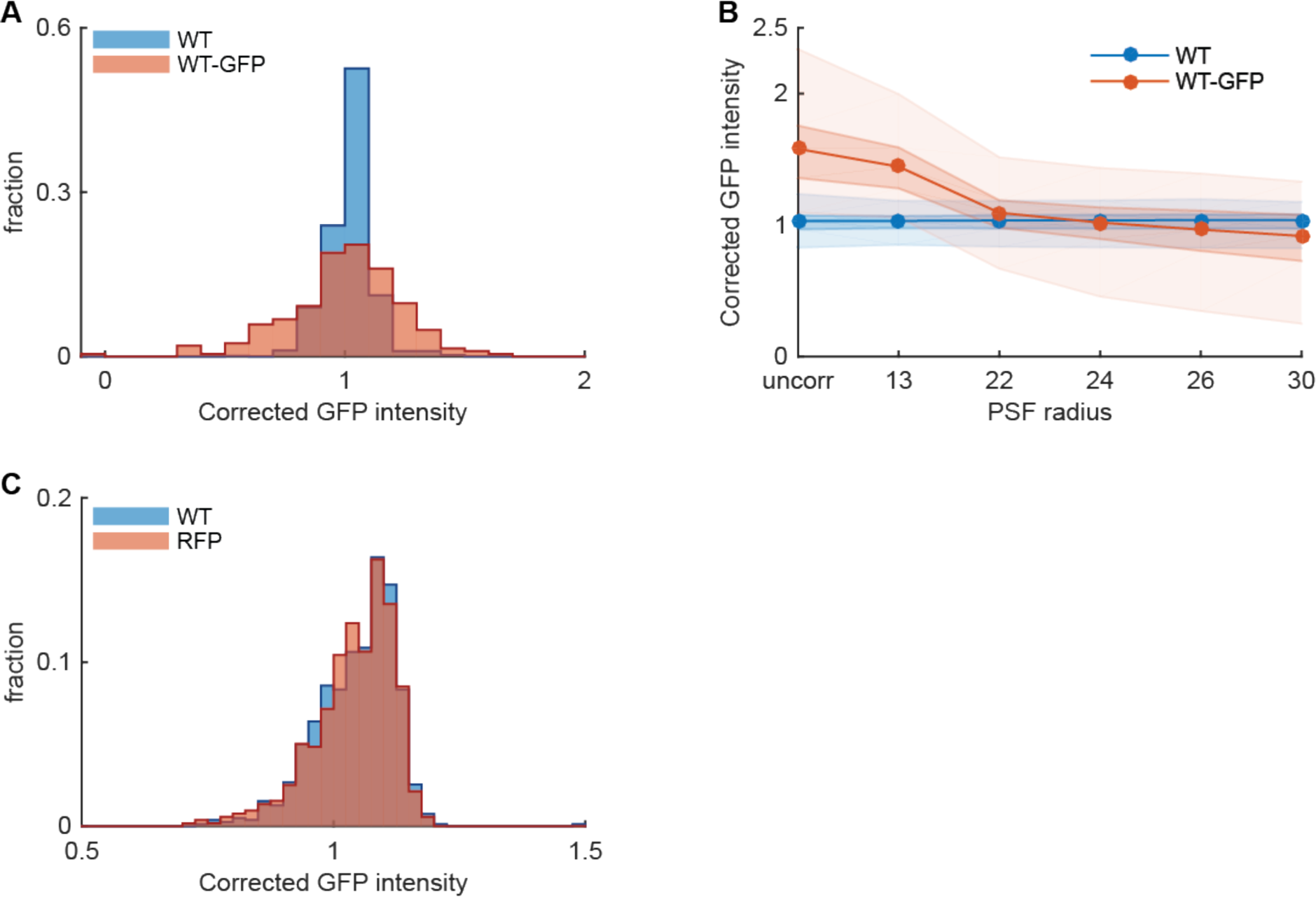
Figure 1–Figure Supplement 3. Correcting for fluorescence halos and bleed-through. **A)** Image correction fully compensates for fluorescence halos. Cells expressing GFP (MG1655+pGFP) were mixed with non-fluorescent wild-type cells (MG1655). There is no significant difference in GFP intensity between isolated wild type cells (WT, blue, median=1.04, interquartile range=(0.97,1.08), n=1118) and wild type cells neighboring a GFP labeled cell (WT-GFP, red, med=1.02, IQR=(0.89,1.13), n=206; p=0.47, Wilcoxon rank sum test). For this figure a point spread function of 24×24 pixels was used (see panel B). **B)** The radius of the point spread function (PSF) strongly affects the accuracy of the deconvolution routine. When the radius is too small (<24) the deconvolution is not strong enough to remove halos from bright cells: i.e. the corrected GFP intensity in WT cells next to GFP labeled cells (red curve) is higher than the intensity in isolated WT cells (blue curve). When the radius of the PSF is too large (>24) artifacts are introduced causing dark halos to appear around bright cells, thus leading to an over correction: i.e. the corrected GFP intensity in WT cells next to GFP labeled cell (red curve) is lower than the intensity in isolated WT cells (blue curve). When a radius of 24 pixels is used, a close to perfect correction is obtained (see panel A). All images are corrected using a PSF of this size. Uncorr: gfp intensities for images that are only background corrected; no shading correction or deconvolution has been performed. Points: median values, dark shaded region: 25-75 percentile range, light shaded region: 2.5-97.5% range. **C)** There is no detectable fluorescence bleed-through from the RFP channel into the GFP channel. Cells expressing RFP (MG1655+pRFP) were mixed with non-fluorescent wild-type cells (MG1655). There is no significant difference in corrected GFP intensities for wild type cells (WT, blue, med=1.06, IQR=(1.00,1.10), n=781) and for RFP labeled cells (RFP, red, med=1.05, IQR=(1.00,1.10), n=517; p=0.51, Wilcoxon rank sum test).

**Figure S4.**
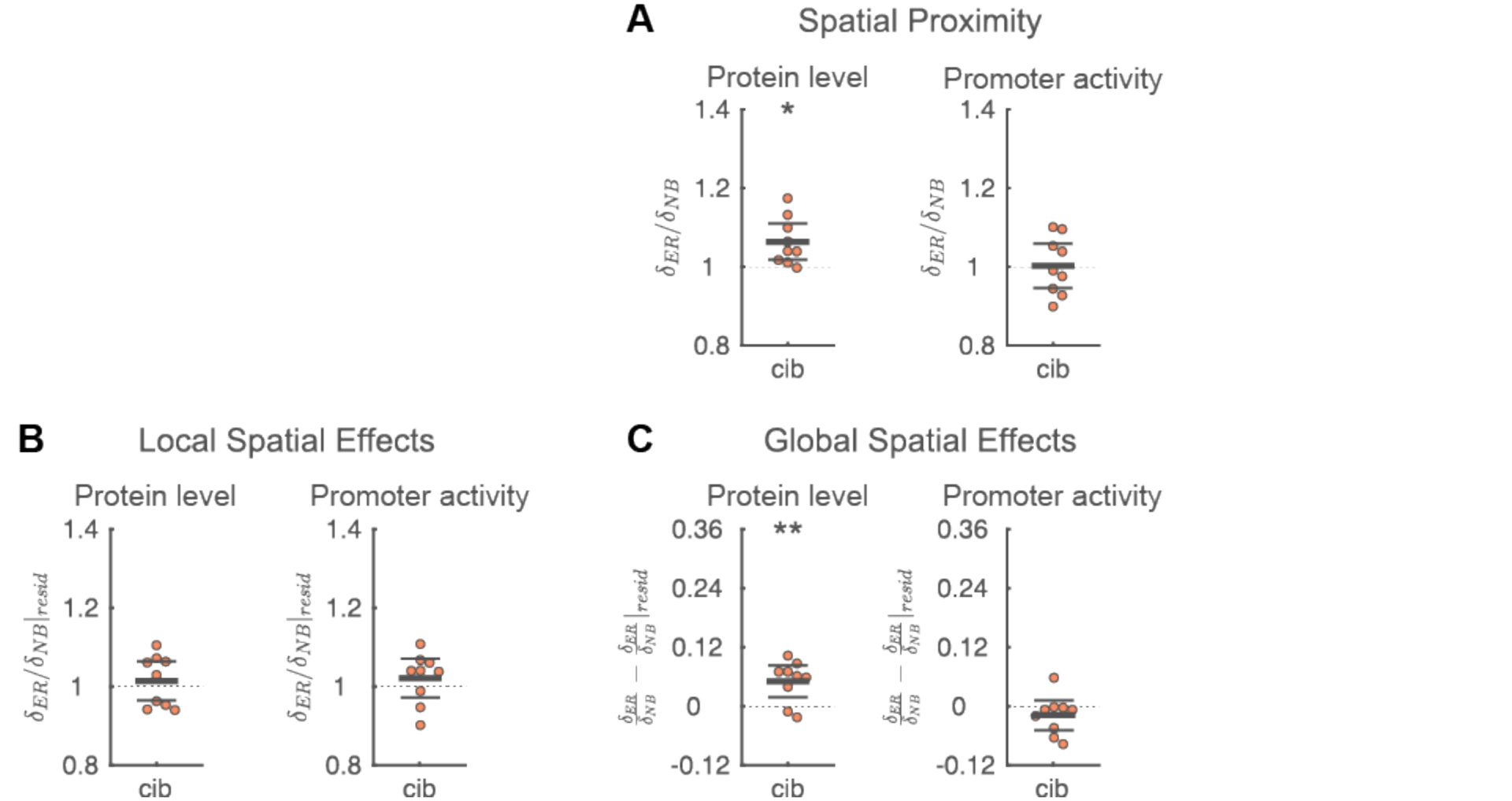
Figure 3–Figure Supplement 1. Robustness of statistic to choice of equally-related cells. Same as Figure 3B-D, except that all equally related cells are considered instead of only the most distant one. Specifically, the median phenotypic distance to all equally related cells is used (see *Methods*). Each point corresponds to a microcolony with 117-138 (mean=128) cells, points are horizontally offset. Thick horizontal lines indicate mean, thin lines 95% confidence intervals. Dashed lines indicate the expected value under the null hypothesis (1 for panel A-B, 0 for panel C). Null hypothesis rejected with: *p<0.05, **p<0.01, ***p<0.001, t-test, *n*=9. Full data can be found in Figure 3 – Figure Supplement 1 – Source Data 1.

**Figure S5.**
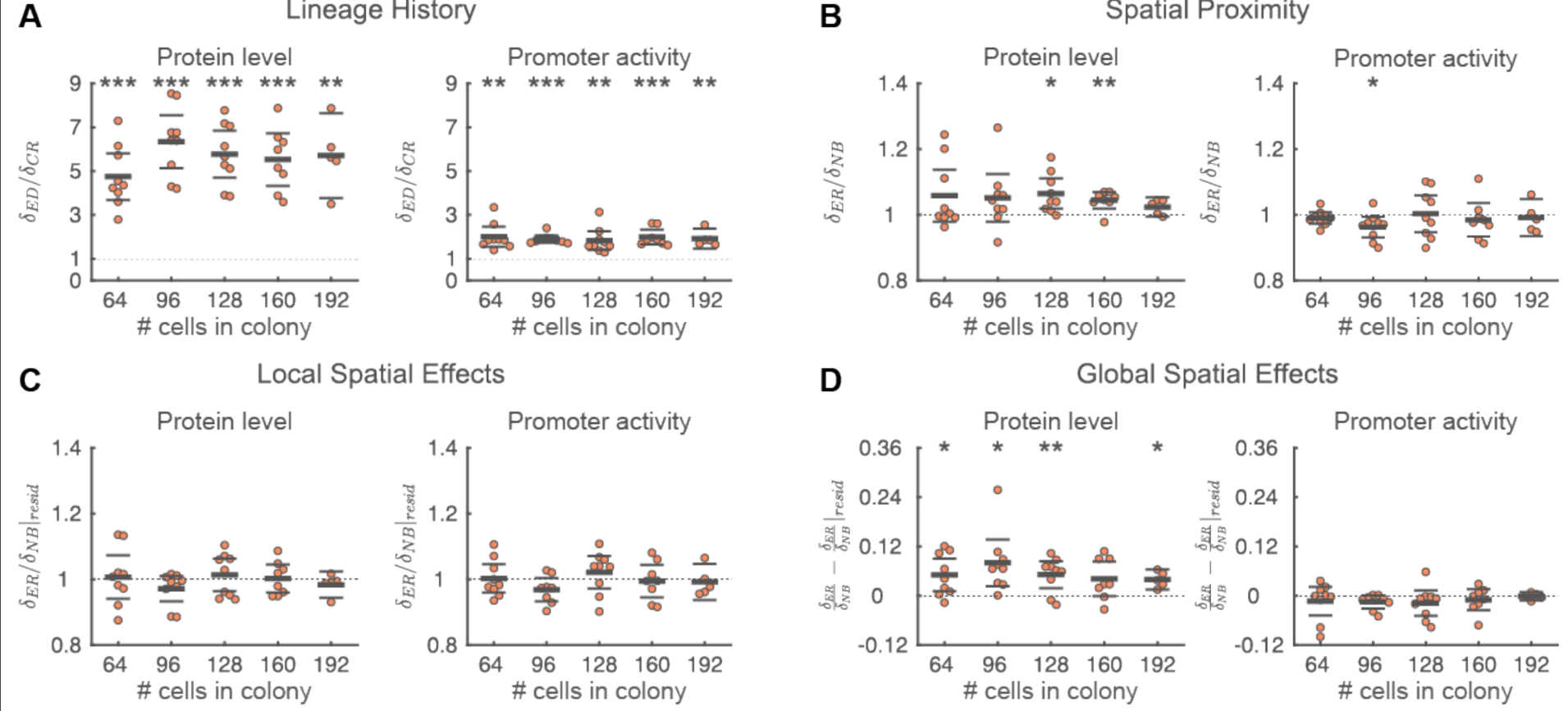
Figure 3–Figure Supplement 2. Robustness of statistic to time of analysis. Same as Figure 3A-D, except that the analysis is repeated for 5 different colony sizes. Each point corresponds to a microcolony of the indicated size, points are horizontally offset. Thick horizontal lines indicate mean, thin lines 95% confidence intervals. Dashed lines indicate the expected value under the null hypothesis (1 for panel A-C, 0 for panel D). Null hypothesis rejected with: *p<0.05, **p<0.01, ***p<0.001, t-test, *n*=9. Full data can be found in Figure 3–Figure Supplement 2 – Source Data 1.

**Figure S6.**
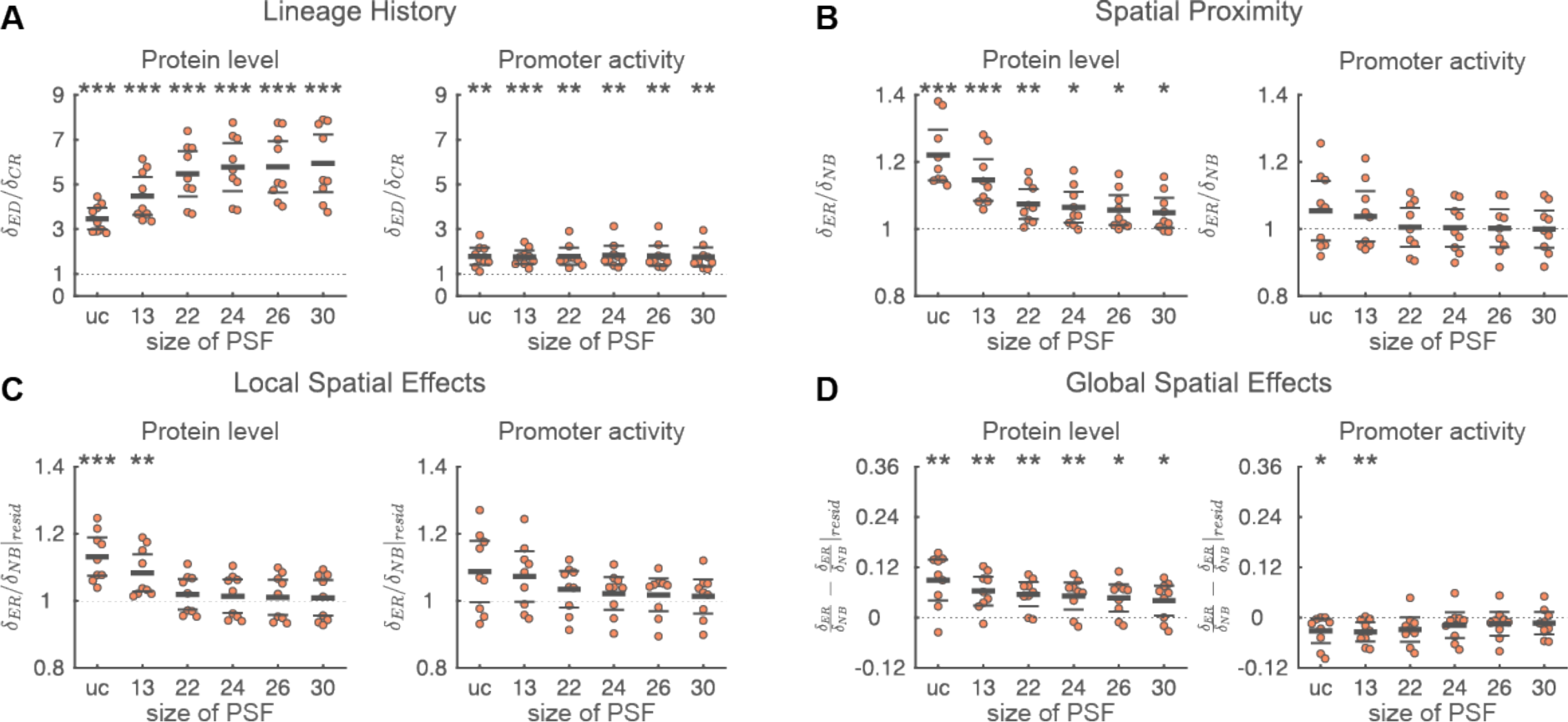
Figure 3–Figure Supplement 3. Robustness of statistic to variation in deconvolution procedure. Same as Figure 3A-D, except that the statistics were calculated for uncorrected intensities (uc) and for deconvoluted data where point spread functions (PSF) of various sizes were used (see also Figure 1–Figure Supplement 3B). PSF radii of 13 (strong under correction), 22 (mild under correction), 24 (optimal correction), 26 (mild over correction) and 30 pixels (strong over correction) were used. The effect of spatial proximity is systematically over estimated for uncorrected and strongly under corrected data (PSF radius=13), but is robust to mild under and over correction (PSF radius between 24 and 30). Uncorrected intensities were calculated from images that are background corrected, but where no shading correction or deconvolution has been applied. Each point corresponds to a microcolony with 117-138 (mean=128) cells, points are horizontally offset. Thick horizontal lines indicate mean, thin lines 95% confidence intervals. Dashed lines indicate the expected value under the null hypothesis (1 for panel A-C, 0 for panel D). Null hypothesis rejected with: *p<0.05, **p<0.01, ***p<0.001, t-test, *n*=9. Full data can be found in Figure 3 – Figure Supplement 3 – Source Data 1.

**Figure S7.**
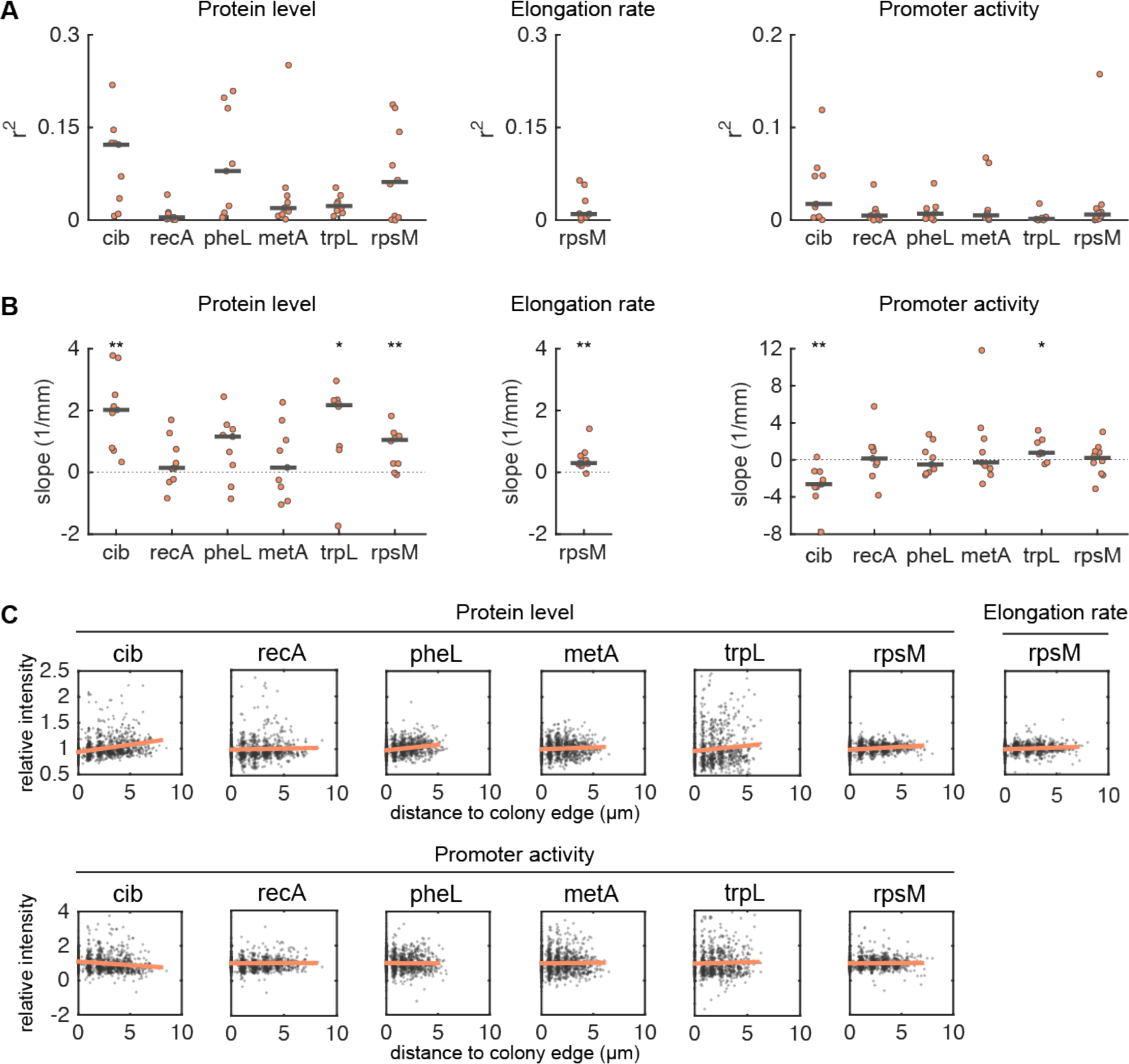
Figure 3–Figure Supplement 4. Dependence of reporter expression dynamics on position in colony. The dependence of protein level, elongation rate, and promotor activity on the position of a cell in the colony. **A)** R-square value of a linear regression of normalized phenotype (i.e. protein level, elongation rate, or promoter activity) with the distance of a cell to the colony edge. Each point corresponds to a microcolony with 117-138 (mean=128) cells, points are horizontally offset. Thick horizontal lines indicate median value. **B)** Slope (1/mm) of a linear regression of normalized phenotype with the distance of a cell to the colony edge. Each point corresponds to a microcolony with 117-138 (mean=128) cells, points are horizontally offset. Thick horizontal lines indicate median value. Dashed lines indicate a slope of 0. Median slope significantly different from zero with: *p<0.05, **p<0.01, Wilcoxon rank sum test, *n* =9, except for *trpL* (*n*=8) and *rpsM* (*n*=10). **C)** Normalized phenotype as function of the distance of a cell to the colony edge. Each point corresponds to a single cell. Data for 8 to 10 microcolonies are pooled together. The red line shows a linear fit to the data. In all cases, normalization is done by dividing the phenotype of a cell by the mean value in the colony.

**Figure S8.**
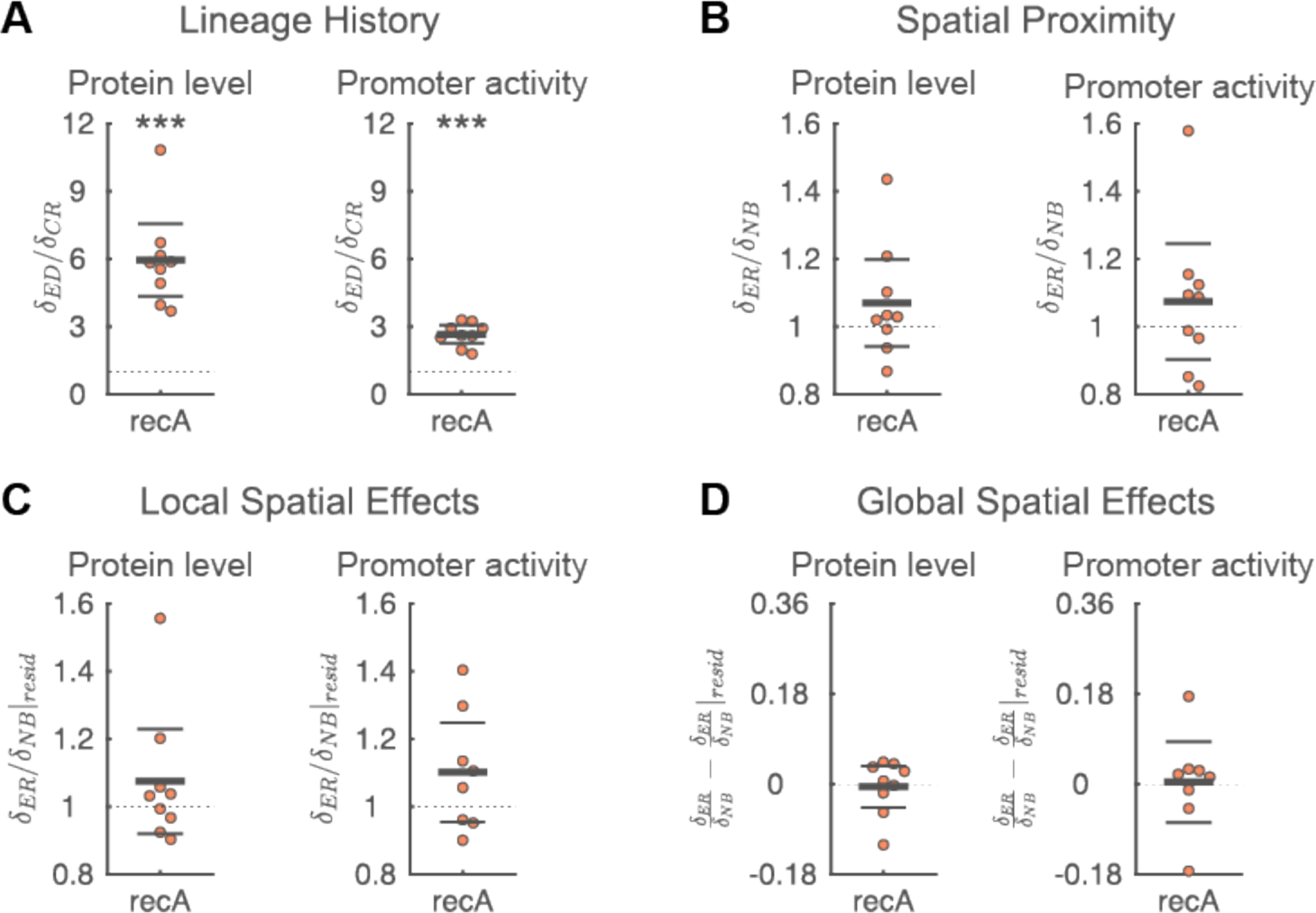
Figure 3–Figure Supplement 5. Expression dynamics of *recA*. Same as Figure 3, but for the expression dynamics of the SOS response gene *recA*. Spatial correlations in the protein level and promoter activity of *recA* are solely due to shared lineage history (panel A), spatial proximity has no significant effect (panel B). Each point corresponds to a microcolony with 117-138 (mean=128) cells, points are horizontally offset. Thick horizontal lines indicate mean, thin lines 95% confidence intervals. Dashed lines indicate the expected value under the null hypothesis (1 for panel A-C, 0 for panel D). Null hypothesis rejected with: *p<0.05, **p<0.01, ***p<0.001, t-test, *n*=9. Full data can be found in Figure 3 – Figure Supplement 5 – Source Data 1.

**Figure S9.**
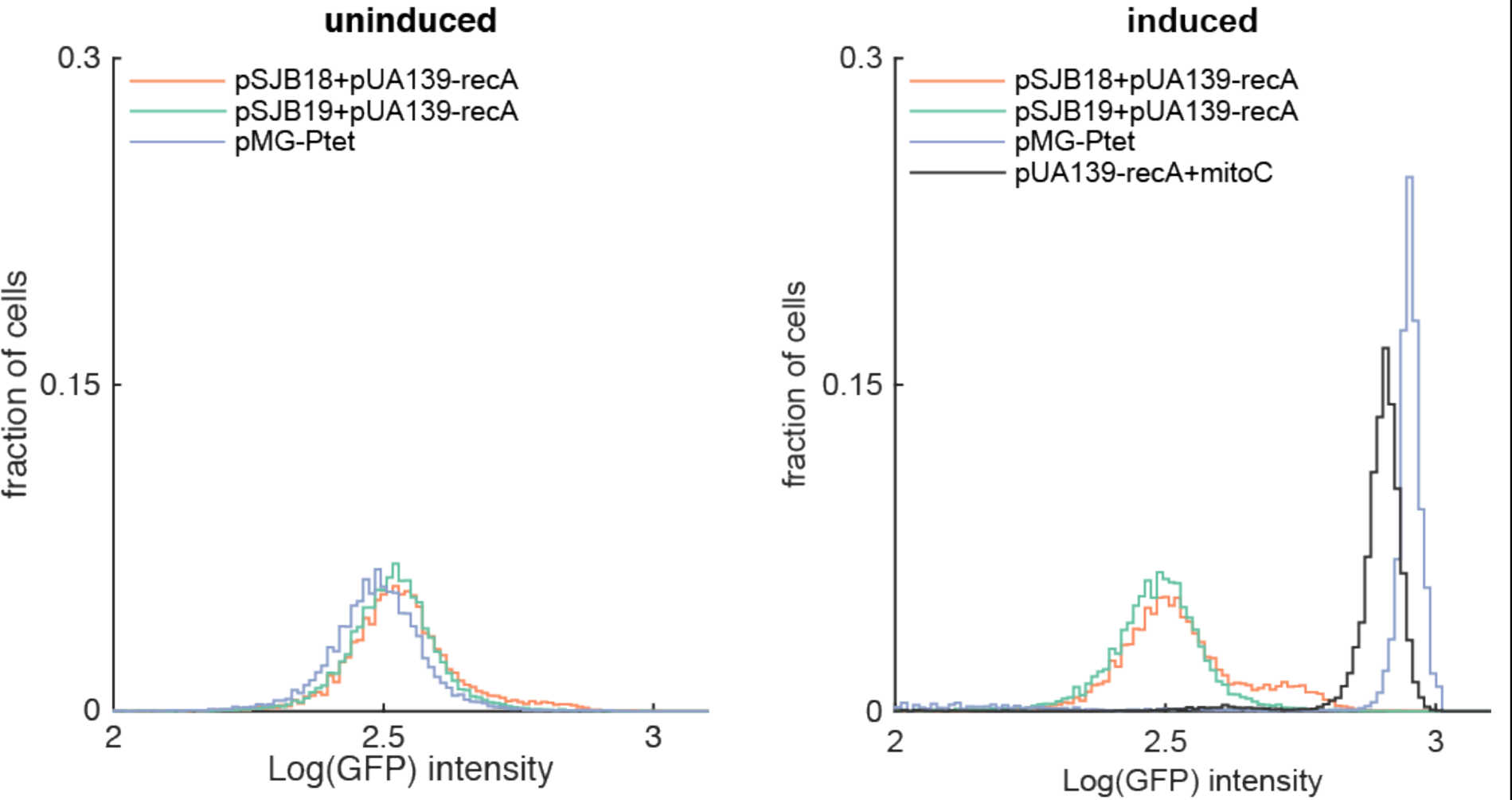
Figure 4–Figure Supplement 1. Inducible SOS response. Histogram of log transformed GFP intensities measured with flow cytometry in uninduced conditions (left) and induced condition (100ng/ml AHT or 0.5µg/ml mitomycin C, right). The expression profiles are shown for 4 strains: **pSJB18+pUA139-recA** (red): containing an inducible nuclease and a GFP transcriptional reporter for *recA*; note that *recA* expression levels increase for a small fraction of cells in the presence of the AHT inducer. **pSJB19+pUA139-recA** (green): same as pSJB18, except that the plasmid also contains the Colicin E2 immunity protein that prevents nuclease activity inside the cell; note that *recA* expression levels do not increase in the presence of AHT. **pMG-pTet** (purple): cell with GFP under control of the P_*tet*_ promoter; note that all cells have increased GFP levels in the presence of AHT. **pUA139-recA** (grey): cell with GFP transcriptional reporter for *recA* where SOS response was induced by adding 0.5µg/ml mitomycin C (mitoC) to the culture medium; note that all cells have high levels of *recA* expression.

**Table 1.** Supplementary File 1–Table 1. **List of plasmids used in this study**. ^†^: pM1437 and p2-camR were co-transformed into MG1655 kanR.

| Name | Carrier strain | Genotype | Description | Source |
| --- | --- | --- | --- | --- |
| **pM1437<sup>†</sup>** | MG1655 kanR (TB60) | pBR322-derivative, *Pcib-gfpmut2*, ampR | *colicin Ib* transcriptional reporter, used together with p2-camR | (Nedialkova et al. 2014) |
| **p2-camR<sup>†</sup>** | MG1655 KanR (TB60) | Salmonella enterica serovar Typhimurium plasmid pColB9, camR | Contains colicin Ib operon, used together with pM1437 | (Stecher et al. 2012) |
| **pUA139-recA** | MG1655 | pUA139 derivative, *PrecA-gfpmut2*, kanR | *recA* transcriptional reporter | (Zaslaver et al. 2006) |
| **pUA139-trpL** | MG1655 | pUA139 derivative, *PtrpL-gfpmut2*, kanR | *trpL* transcriptional reporter | (Zaslaver et al. 2006) |
| **pUA139-rpsM** | MG1655 | pUA139 derivative, *PrpsM-gfpmut2*, kanR | *rpsM* transcriptional reporter | (Zaslaver et al. 2006) |
| **pUA66-metA** | MG1655 | pUA66 derivative, *PmetA-gfpmut2*, kanR | *metA* transcriptional reporter | (Zaslaver et al. 2006) |
| **pUA66-pheA** | MG1655 | pUA66 derivative, *PpheA-gfpmut2*, kanR | *pheA* transcriptional reporter | (Zaslaver et al. 2006) |
| **pSV66-recA-rpsM** | MG1655 | pUA139 derivative, *PrecA-gfpmut2*, *PrpsM-turboRFP*, kanR | *recA/rpsM* dual transcriptional reporter | This study |
| **pGFP** | MG1655 | pQE derivative, *Plac-gfpmut2*, ampR | IPTG inducible *gfp* | Lab strain-collection |
| **pRFP** | MG1655 | pQE derivative, *Plac-mcherry*, ampR | IPTG inducible *rfp* | Lab strain-collection |
| **pSJB18** | MG1655 nalR smR | pMG-Ptet derivative, *Ptet-cea*(nuclease domain), ampR | Tet inducible nuclease domain of colicin E2 | This study |
| **pSJB19** | MG1655 nalR smR | pMG-Ptet derivative, *Ptet-cea*(nuclease domain)-*cei*, ampR | Tet inducible nuclease domain of colicin E2 + colicin E2 immunity protein | This study |
| **pMG-pTet** | MG1655 nalR smR | *Ptet-gfp* | Tet inducible GFP | (Neuenschwander et al. 2007) |
| **pTurboRFP** | MG1655 | *turboRFP* | Source of turboRFP coding sequence | (Hol et al. 2014) |
| **pZS2-123** | MG1655 | Triple reporter | Source of multiple terminator sequence | (Cox, Dunlop, and Elowitz 2010) |
| **pColE2-p9** | MG1655 | Colicin E2 plasmid | Source of *cea* and *cei* coding sequence | (Hol et al. 2014) |

## Supplementary File 2 – Plasmid construction

### Construction of pSJB18 and pSJB19

The pSJB18 plasmid is based on the pMG-Ptet plasmid (Neuenschwander et al. 2007), Neuenschwander et al. 2007 which contains the tetracycline inducible P_tet_ promoter. The *colicin E2* nuclease domain (*cea*_*nuclease*_) was amplified from plasmid pColE2-p9 (Kerr et al. 2002; Hol et al. 2014) using primers ColE2_C-dom_tet_fwd and ColE2_tet _rev (see Supplementary File 2–Table 1 for primer sequences). The forward primer includes an ATG start codon and Xbal restriction site, the reverse primer includes a Xhol restriction site. The amplicon was inserted into pMG-P_tet_ via Xbal/Xhol. The construct sequence was confirmed with sequencing.

We constructed a second plasmid, pSJB19, that is identical to pSJB18, except that it also includes the Colicin E2 immunity protein (*cei*). This immunity protein is co-expressed with the nuclease domain and inhibits the nuclease activity. The Colicin E2 nuclease domain ( *cea*_*nuclease*_) was amplified together with the colicin E2 immunity protein (*cei*) from plasmid pColE2-p9 (Kerr et al. 2002; Hol et al. 2014) using primers ColE2_C-dom_tet_fwd and ImmE2_tet_rev (see Supplementary File 2–Table 1). The forward primer includes an ATG start codon and Xbal restriction site, the reverse primer includes a Xhol restriction site. The amplicon was inserted into pMG-P_tet_ via Xbal/Xhol. The construct sequence was confirmed with sequencing.

### Construction of pSV66-rpsM-rpsM dual-reporter

The pSV66 dual reporter is based on the low copy number pUA66/pUA139 (Zaslaver et al. 2006) reporter plasmid, but contains an additional transcriptional reporter based on the red fluorescent protein TurboRFP. The arrangement of the two reporters was based on the triple reporter plasmid pZS2-123 (Cox, Dunlop, and Elowitz 2010). Specifically, the *gfmut2* and *turborfp* reporters are separated by a region containing multiple transcriptional terminators amplified from pZS2-123. The plasmid was constructed using a three-step Gibson assembly protocol (New England Biolabs, Ipswich, Massachusetts):

1. *1) PrpsM-tRFP* promoter-reporter construct. The promotor region of *rpsM* was amplified from plasmid pUA139-rpsM (Zaslaver et al. 2006) using primers Prpsm-fw/rv (see Supplementary File 2–Table 1 for primer sequences). The coding sequence of *turborfp* was amplified from plasmid pTurboRFP (Hol et al. 2014) using rfp-fw/rv. The resulting products were joined using Gibson assembly.
2. *2) PrpsM-turboRFP-terminator* construct. The *PrpsM-turboRFP* construct from step 1 was amplified from the Gibson assembly product using primers Prpsm-fw and rfp-rv. The multiple terminator region was amplified from plasmid pZS2-123 (Cox, Dunlop, and Elowitz 2010) using primers ter-fw/rv. The resulting products were joined using Gibson assembly.
3. 3) pSV66-rpsM-rpsM plasmid. The *PrpsM-turboRFP-terminator* construct from step 2 was amplified from the Gibson assembly product using primers ter-rv and Prfp-fw. The greater part of plasmid pUA139-rpsM ((Zaslaver et al. 2006), including the *Prpsm-gfpmut2* reporter, origin of replication and kanamycin resistance cassette) was amplified using primers vector-fw/rv. The resulting products were joined using Gibson assembly.

We confirmed the sequence of the promoter regions, fluorescent proteins, terminator region and origin of replication using sequencing.

### Construction of pSV66-recA-rpsM dual-reporter

To construct the dual reporter pSV66-recA-rpsM the promoter region in front of *gfpmut2* of the pSV66-rpsM-rpsM was replaced with the promoter of *recA* using a one step Gibson assembly process. The promoter for *recA* was amplified from the plasmid pUA139-recA (Zaslaver et al. 2006) using primer Pgfp-fw/rv and the backbone of pSV66-rpsM-rpsM was amplified with gfpVec-fw/rv. The two PCR products were then combined using Gibson assembly and the sequence of both the *recA* and *rpsM* promoter region, as well as the intermediate terminator region was confirmed using sequencing.

In all steps we used the Q5 high-fidelity DNA polymerase for DNA amplification and Gibson Assembly master mix for Gibson assembly (New England Biolabs, Ipswich, Massachusetts).

## References

Ackermann, Martin. 2015. “A Functional Perspective on Phenotypic Heterogeneity in Microorganisms.” Nature Reviews Microbiology 13 (8). Nature Publishing Group: 497–508. doi:10.1038/nrmicro3491.

Ackermann, Martin, Bärbel Stecher, Nikki E Freed, Pascal Songhet, Wolf-Dietrich Hardt and Michael Doebeli. 2008. “Self-Destructive Cooperation Mediated by Phenotypic Noise.” Nature 454 (7207): 987–90. doi:10.1038/nature07067.

Bassler, Bonnie L, and Richard Losick. 2006. “Bacterially Speaking.” Cell 125 (2): 237–46. doi:10.1016/j.cell.2006.04.001.

Berdahl, Andrew, Colin J Torney, Christos C Ioannou, Jolyon J Faria, and Iain D Couzin. 2013. “Emergent Sensing of Complex Environments by Mobile Animal Groups.” Science 339 (6119). American Association for the Advancement of Science: 574–76. doi:10.1126/science.1225883.

Cascales, Eric, Susan K Buchanan, Denis Duché, Colin Kleanthous, Roland Lloubès, Kathleen Postle, Margaret Riley, Stephen Slatin and Danièle Cavard. 2007. “Colicin Biology.” Microbiology and Molecular Biology Reviews: MMBR 71 (1). American Society for Microbiology: 158–229. doi:10.1128/MMBR.00036-06.

Chubukov, Victor, Luca Gerosa, Karl Kochanowski, and Uwe Sauer. 2014. “Coordination of Microbial Metabolism.” Nature Reviews Microbiology 12 (5). Nature Publishing Group: 327–40. doi:10.1038/nrmicro3238.

Claessen, Dennis, Daniel E Rozen, Oscar P Kuipers, Lotte Søgaard-Andersen, and Gilles P van Wezel. 2014. “Bacterial Solutions to Multicellularity: a Tale of Biofilms, Filaments and Fruiting Bodies.” Nature ReviewsMicrobiology 12 (2): 115–24. doi:10.1038/nrmicro3178.

Dubey, Gyanendra P, and Sigal Ben-Yehuda. 2011. “Intercellular Nanotubes Mediate Bacterial Communication.” Cell 144 (4): 590–600. doi:10.1016/j.cell.2011.01.015.

Ducret, Adrien, Betty Fleuchot, Ptissam Bergam, Tâm Mignot and Peter Greenberg. 2013. “Direct Live Imaging of Cell–Cell Protein Transfer by Transient Outer Membrane Fusion in Myxococcus Xanthus.” eLife 2 (July). eLife Sciences Publications Limited: e00868. doi:10.7554/eLife.00868.

Elowitz, Michael B, Arnold J Levine, Eric D Siggia, and Peter S Swain. 2002. “Stochastic Gene Expression in a Single Cell.” 297 (5584): 1183–86. doi:10.1126/science.1070919.

Flemming, Hans-Curt, Jost Wingender, Ulrich Szewzyk, Peter Steinberg, Scott A Rice, and Staffan Kjelleberg. 2016. “Biofilms: an Emergent Form of Bacterial Life.” Nature Reviews Microbiology 14 (9). Nature Research: 563–75. doi:10.1038/nrmicro.2016.94.

Friedman, Nir, Shuki Vardi, Michal Ronen, Uri Alon and Joel Stavans. 2005. “Precise Temporal Modulation in the Response of the SOS DNA Repair Network in Individual Bacteria.” PLoS Biology 3 (7). Public Library of Science: e238. doi:10.1371/journal.pbio.0030238.

Guantes, Raúl, Ilaria Benedetti, Rafael Silva-Rocha, and Víctor de Lorenzo. 2015. “Transcription Factor Levels Enable Metabolic Diversification of Single Cells of Environmental Bacteria.” The ISME Journal 10 (5): 1122–33. doi:10.1038/ismej.2015.193.

Hayes, Christopher S, Stephanie K Aoki, and David A Low. 2010. “Bacterial Contact-Dependent Delivery Systems.” Dx.Doi.org 44 (1). Annual Reviews: 71–90. doi:10.1146/annurev.genet.42.110807.091449.

Hein, Andrew M, Sara Brin Rosenthal, George I Hagstrom, Andrew Berdahl, Colin J Torney and Iain D Couzin. 2015. “The Evolution of Distributed Sensing and Collective Computation in Animal Populations.” eLife 4 (December): e10955. doi:10.7554/eLife.10955.

Hormoz, Sahand, Nicolas Desprat, and Boris I Shraiman. 2015. “Inferring Epigenetic Dynamics From Kin Correlations.” Proc Natl Acad Sci USA 112 (18). National Acad Sciences: E2281–89. doi:10.1073/pnas.1504407112.

Johnson, David R, Felix Goldschmidt, Elin E Lilja, and Martin Ackermann. 2012. “Metabolic Specialization and the Assembly of Microbial Communities.” The ISME Journal, May. doi:10.1038/ismej.2012.46.

Julou, T, T Mora, L Guillon, V Croquette, I J Schalk, D Bensimon, and N Desprat. 2013. “Cell-Cell Contacts Confine Public Goods Diffusion Inside Pseudomonas Aeruginosa Clonal Microcolonies.” Proceedings of the National Academy of Sciences of the United States of America 110 (31): 12577–82. doi:10.1073/pnas.1301428110.

Kaern, Mads, Timothy C Elston, William J Blake, and James J Collins. 2005. “Stochasticity in Gene Expression: From Theories to Phenotypes.” Nature Reviews Genetics 6 (6): 451–64. doi:10.1038/nrg1615.

Kiviet, Daniel J, Philippe Nghe, Noreen Walker, Sarah Boulineau, Vanda Sunderlikova and Sander J Tans. 2014. “Stochasticity of Metabolism and Growth at the Single-Cell Level.” Nature 514 (7522): 376–79. doi:10.1038/nature13582.

Kochanowski, Karl, Uwe Sauer, and Elad Noor. 2015. “Posttranslational Regulation of Microbial Metabolism.” Current Opinion in Microbiology 27 (October): 10–17. doi:10.1016/j.mib.2015.05.007.

Kolter, Roberto, Hera Vlamakis, and Jordi van Gestel. 2015. “Division of Labor in Biofilms: the Ecology of Cell Differentiation. - PubMed - NCBI.” Microbiology Spectrum 3 (2): MB–0002–2014. doi:10.1128/microbiolspec.MB-0002-2014.

Lee, Heewook, Ellen Popodi, Haixu Tang, and Patricia L Foster. 2012. “Rate and Molecular Spectrum of Spontaneous Mutations in the Bacterium Escherichia Coli as Determined by Whole-Genome Sequencing.” Proc Natl Acad Sci USA 109 (41): E2774–83. doi:10.1073/pnas.1210309109.

Liu, Wenzheng, Henriette L Røder, Jonas S Madsen, Thomas Bjarnsholt, Søren J Sørensen and Mette Burmølle. 2016. “Interspecific Bacterial Interactions Are Reflected in Multispecies Biofilm Spatial Organization.” Name: Frontiers in Microbiology 7 (258): 4258. doi:10.3389/fmicb.2016.01366.

Lopez, Daniel, and Roberto Kolter. 2010. “Extracellular Signals That Define Distinct and Coexisting Cell Fates in Bacillus Subtilis.” FEMS Microbiology Reviews 34 (2): 134–49. doi:10.1111/j.1574-6976.2009.00199.x.

Mee, Michael T, James J Collins, George M Church, and Harris H Wang. 2014. “Syntrophic Exchange in Synthetic Microbial Communities.” Proc Natl Acad Sci USA 111 (20). National Acad Sciences: E2149–56. doi:10.1073/pnas.1405641111.

Muro-Pastor, Alicia M, and Wolfgang R Hess. 2012. “Heterocyst Differentiation: From Single Mutants to Global Approaches.” Trends in Microbiology 20 (11): 548–57. doi:10.1016/j.tim.2012.07.005.

Nadell, C D, V Bucci, K Drescher, S A Levin, B L Bassler, and J B Xavier. 2013. “Cutting Through the Complexity of Cell Collectives.” Proceedings. Biological Sciences / the RoyalSociety 280 (1755): 20122770–70. doi:10.1098/rspb.2012.2770.

Nadell, Carey D, Knut Drescher, and Kevin R Foster. 2016. “Spatial Structure, Cooperation and Competition in Biofilms.” Nature Reviews Microbiology 14 (9). Nature Research: 589–600. doi:10.1038/nrmicro.2016.84.

Nedialkova, Lubov Petkova, Rémy Denzler, Martin B Koeppel, Manuel Diehl, Diana Ring, Thorsten Wille, Roman G Gerlach and Bärbel Stecher. 2014a. “Inflammation Fuels Colicin Ib-Dependent Competition of Salmonella Serovar Typhimurium and E. Coli in Enterobacterial Blooms.” Edited by Jorge E Galán. PLoS Pathogens 10 (1). Public Library of Science: e1003844. doi:10.1371/journal.ppat.1003844.

Nedialkova, Lubov Petkova, Rémy Denzler, Martin B Koeppel, Manuel Diehl, Diana Ring, Thorsten Wille, Roman G Gerlach and Bärbel Stecher. 2014b. “Inflammation Fuels Colicin Ib-Dependent Competition of Salmonella Serovar Typhimurium and E. Coli in Enterobacterial Blooms.” Edited by Jorge E Galán. PLoS Pathogens 10 (1). Public Library of Science: e1003844. doi:10.1371/journal.ppat.1003844.

Ozbudak, Ertugrul M, Mukund Thattai, Iren Kurtser, Alan D Grossman and Alexander van Oudenaarden. 2002. “Regulation of Noise in the Expression of a Single Gene.” Nature Genetics 31 (1): 69–73. doi:10.1038/ng869.

Pande, Samay, Holger Merker, Katrin Bohl, Michael Reichelt, Stefan Schuster, Luís F de Figueiredo, Christoph Kaleta, and Christian Kost. 2013. “Fitness and Stability of Obligate Cross-Feeding Interactions That Emerge Upon Gene Loss in Bacteria.” The ISME Journal, November. doi:10.1038/ismej.2013.211.

Pande, Samay, Shraddha Shitut, Lisa Freund, Martin Westermann, Felix Bertels, Claudia Colesie, Ilka B Bischofs and Christian Kost. 2015. “Metabolic Cross-Feeding via Intercellular Nanotubes Among Bacteria.” Nature Communications 6 (February). Nature Publishing Group: 6238. doi:10.1038/ncomms7238.

Pennington, Jeanine M, and Susan M Rosenberg. 2007. “Spontaneous DNA Breakage in Single Living Escherichia Coli Cells.” Nature Genetics 39 (6): 797–802. doi:10.1038/ng2051.

Popat, R, D M Cornforth, L McNally, and S P Brown. 2014. “Collective Sensing and Collective Responses in Quorum-Sensing Bacteria.” Journal of the Royal Society Interface 12 (103): 20140882–82. doi:10.1098/rsif.2014.0882.

Riley, Margaret A, and John E Wertz. 2002. “Bacteriocins: Evolution, Ecology, and Application.” Annual Review of Microbiology 56 (1): 117–37. doi:10.1146/annurev.micro.56.012302.161024.

Risser, Douglas D, Francis C Y Wong, and John C Meeks. 2012. “Biased Inheritance of the Protein PatN Frees Vegetative Cells to Initiate Patterned Heterocyst Differentiation.” Proc Natl Acad Sci USA 109 (38). National Acad Sciences: 15342–47. doi:10.1073/pnas.1207530109.

Robert, Lydia, Gregory Paul, Yong Chen, François Taddei, Damien Baigl and Ariel B Lindner. 2010. “Pre-Dispositions and Epigenetic Inheritance in the Escherichia Coli Lactose Operon Bistable Switch.” Molecular Systems Biology 6 (April): 357. doi:10.1038/msb.2010.12.

Ross-Gillespie, Adin, and Rolf Kümmerli. 2014. “Collective Decision-Making in Microbes.” Name: Frontiers in Microbiology 5. doi:10.3389/fmicb.2014.00054.

Schindelin, Johannes, Ignacio Arganda -Carreras, Erwin Frise, Verena Kaynig, Mark Longair, Tobias Pietzsch, Stephan Preibisch, et al. 2012. “Fiji: an Open-Source Platform for Biological-Image Analysis.” Nature Methods 9 (7): 676–82. doi:10.1038/nmeth.2019.

Schwechheimer, Carmen, and Meta J Kuehn. 2015. “Outer-Membrane Vesicles From Gram-Negative Bacteria: Biogenesis and Functions.” Nature Reviews Microbiology 13 (10). Nature Research: 605–19. doi:10.1038/nrmicro3525.

Scott, M, S Klumpp, E M Mateescu, and T Hwa. 2014. “Emergence of Robust Growth Laws From Optimal Regulation of Ribosome Synthesis.” Molecular Systems Biology 10 (8): 747–47. doi:10.15252/msb.20145379.

Scott, Matthew, Carl W Gunderson, Eduard M Mateescu, Zhongge Zhang, and Terence Hwa. 2010. “Interdependence of Cell Growth and Gene Expression: Origins and Consequences.” Science 330 (6007): 1099–1102. doi:10.1126/science.1192588.

Silander, Olin K, Nela Nikolic, Alon Zaslaver, Anat Bren, Ilya Kikoin, Uri Alon and Martin Ackermann. 2012. “A Genome-Wide Analysis of Promoter-Mediated Phenotypic Noise in Escherichia Coli.” Plos Genetics 8 (1): e1002443. doi:10.1371/journal.pgen.1002443.

Simons, A. 2004. “Many Wrongs: the Advantage of Group Navigation.” Trends in Ecology & Evolution 19 (9): 453–55. doi:10.1016/j.tree.2004.07.001.

Snijder, Berend, and Lucas Pelkmans. 2011. “Origins of Regulated Cell-to-Cell Variability.” Nature Reviews Molecular Cell Biology 12 (2): 119–25. doi:10.1038/nrm3044.

Spriewald, Stefanie, Jana Glaser, Markus Beutler, Martin B Koeppel and Bärbel Stecher. 2015. “Reporters for Single-Cell Analysis of Colicin Ib Expression in Salmonella Enterica Serovar Typhimurium.” Edited by Eric Cascales. PLoS ONE 10 (12): e0144647. doi:10.1371/journal.pone.0144647.

Stecher, Bärbel, Rémy Denzler, Lisa Maier, Florian Bernet, Mandy J Sanders, Derek J Pickard, Manja Barthel, et al. 2012. “Gut Inflammation Can Boost Horizontal Gene Transfer Between Pathogenic and Commensal Enterobacteriaceae.” Proc Natl Acad Sci USA 109 (4): 1269–74. doi:10.1073/pnas.1113246109.

Stewart, Philip S, and Michael J Franklin. 2008. “Physiological Heterogeneity in Biofilms.” Nature Reviews Microbiology 6 (3). Nature Publishing Group: 199–210. doi:10.1038/nrmicro1838.

Symmons, Orsolya, and Arjun Raj. 2016. “What’s Luck Got to Do with It: Single Cells, Multiple Fates, and Biological Nondeterminism.” Molecular Cell 62 (5). Elsevier: 788–802. doi:10.1016/j.molcel.2016.05.023.

van Gestel, Jordi, Hera Vlamakis, and Roberto Kolter. 2015. “From Cell Differentiation to Cell Collectives: Bacillus Subtilis Uses Division of Labor to Migrate.” Edited by Michael T Laub. PLoS Biology 13 (4). Public Library of Science: e1002141. doi:10.1371/journal.pbio.1002141.

van Vliet, Simon, and Martin Ackermann. 2015. “Bacterial Ventures Into Multicellularity: Collectivism Through Individuality.” PLoS Biology 13 (6). Public Library of Science: e1002162. doi:10.1371/journal.pbio.1002162.

Veening, J W, E J Stewart, T W Berngruber, F Taddei, O P Kuipers, and L W Hamoen. 2008. “Bet-Hedging and Epigenetic Inheritance in Bacterial Cell Development.” Proceedings of the National Academy of Sciences of the United States of America 105 (11): 4393–98. doi:10.1073/pnas.0700463105.

Veening, Jan-Willem, Wiep Klaas Smits, and Oscar P Kuipers. 2008. “Bistability, Epigenetics, and Bet-Hedging in Bacteria.” Annual Review of Microbiology 62 (1): 193–210. doi:10.1146/annurev.micro.62.081307.163002.

Wintermute, Edwin H, and Pamela A Silver. 2010. “Emergent Cooperation in Microbial Metabolism.” Molecular Systems Biology 6 (September): 407. doi:10.1038/msb.2010.66.

Young, Jonathan W, James C W Locke, Alphan Altinok, Nitzan Rosenfeld, Tigran Bacarian, Peter S Swain, Eric Mjolsness, and Michael B Elowitz. 2011. “Measuring Single-Cell Gene Expression Dynamics in Bacteria Using Fluorescence Time-Lapse Microscopy.” Nature Protocols 7 (1): 80–88. doi:10.1038/nprot.2011.432.

Zaslaver, Alon, Anat Bren, Michal Ronen, Shalev Itzkovitz, Ilya Kikoin, Seagull Shavit, Wolfram Liebermeister, Michael G Surette and Uri Alon. 2006. “A Comprehensive Library of Fluorescent Transcriptional Reporters for Escherichia Coli.” Nature Methods 3 (8): 623–28. doi:10.1038/nmeth895.

## References – Supplementary File 1

Cox, Robert S, Mary J Dunlop, and Michael B Elowitz. 2010. “A Synthetic Three-Color Scaffold for Monitoring Genetic Regulation and Noise.” Journal of Biological Engineering 4 (1). BioMed Central Ltd: 10. doi:10.1186/1754-1611-4-10.

Hol, Felix, Mathias J Voges, Cees Dekker, and Juan E Keymer. 2014. “Nutrient-Responsive Regulation Determines Biodiversity in a Colicin-Mediated Bacterial Community.” BMC Biology 12 (1): 68. doi:10.1186/s12915-014-0068-2.

Nedialkova, Lubov Petkova, Rémy Denzler, Martin B Koeppel, Manuel Diehl, Diana Ring, Thorsten Wille, Roman G Gerlach and Bärbel Stecher. 2014. “Inflammation Fuels Colicin Ib-Dependent Competition of Salmonella Serovar Typhimurium and E. Coli in Enterobacterial Blooms.” Edited by Jorge E Galán. PLoS Pathogens 10 (1). Public Library of Science: e1003844. doi:10.1371/journal.ppat.1003844.

Neuenschwander, Martin, Maren Butz, Caroline Heintz, Peter Kast and Donald Hilvert. 2007. “A Simple Selection Strategy for Evolving Highly Efficient Enzymes.” Nature Biotechnology 25 (10): 1145–47. doi:10.1038/nbt1341.

## References – Supplementary File 2

Kerr, Benjamin, Margaret A Riley, Marcus W Feldman, and Brendan J M Bohannan. 2002. “Local Dispersal Promotes Biodiversity in a Real-Life Game of Rock-Paper-Scissors.” Nature 418 (6894). Nature Publishing Group: 171–74. doi:10.1038/nature00823.

